# Endogenous retrovirus-like proteins recruit UBQLN2 to stress granules and alter their functional properties

**DOI:** 10.1101/2024.10.24.620053

**Authors:** Harihar M. Mohan, Martin G. Fernandez, Camellia Huang, Rita Lin, Jaimie H. Ryou, Donald Seyfried, Nikolas Grotewold, Alexandra M. Whiteley, Sami J. Barmada, Venkatesha Basrur, Shyamal Mosalaganti, Henry L. Paulson, Lisa M. Sharkey

**Affiliations:** Cellular and Molecular Biology Program, University of Michigan Medical School, Ann Arbor, MI 48109, United States; Department of Neurology, University of Michigan Medical School, Ann Arbor, MI 48109, United States; Life Sciences Institute, University of Michigan, Ann Arbor, MI 48109, United States; Department of Biophysics, College of Literature, Science and the Arts, University of Michigan, 48109, United States; Medical Scientist Training Program, University of Michigan Medical School, Ann Arbor, MI 48109, United States; Department of Biochemistry, University of Colorado Boulder, Boulder, CO 80309, United States; University of Michigan Proteomics Resource Facility, University of Michigan Medical School, Ann Arbor, MI 48109, United States; Department of Cell and Developmental Biology, University of Michigan Medical School, Ann Arbor, MI 48109, United States; Department of Biological Chemistry, University of Michigan Medical School, Ann Arbor, MI 48109, United States

## Abstract

The human genome is replete with sequences derived from foreign elements including endogenous retrovirus-like proteins of unknown function. Here we show that UBQLN2, a ubiquitin-proteasome shuttle factor implicated in neurodegenerative diseases, is regulated by the linked actions of two retrovirus-like proteins, RTL8 and PEG10. RTL8 confers on UBQLN2 the ability to complex with and regulate PEG10. PEG10, a core component of stress granules, drives the recruitment of UBQLN2 to stress granules under various stress conditions, but can only do so when RTL8 is present. Changes in PEG10 levels further remodel the kinetics of stress granule disassembly and overall composition by incorporating select extracellular vesicle proteins. Within stress granules, PEG10 forms virus-like particles, underscoring the structural heterogeneity of this class of biomolecular condensates. Together, these results reveal an unexpected link between pathways of cellular proteostasis and endogenous retrovirus-like proteins.

## INTRODUCTION

Nearly 8% of the human genome is derived from long terminal repeat (LTR)-containing retroelements such as retroviruses and retrotransposons.^1,2^ While most LTR retroelements have lost their functionality or are silenced,^3^ some have been repurposed to encode protein-coding genes that serve important functions in physiology and disease. Prominent examples include the activity-regulated cytoskeleton-associated protein (Arc), the paraneoplastic Ma antigen (PNMA) family, and the Retrotransposon Gag-like (RTL) family, each of which is derived from a unique ancestor.^4^ Arc is critical for synaptic plasticity, and the less characterized PNMA and RTL families have been implicated in development, innate immunity, cancer, and neonatal syndromes.^5–18^ Two members of the RTL family, paternally expressed gene 10 (PEG10) and RTL8, are linked to neurological diseases and malignancies yet their biological functions remain uncertain.^15,19–26^ Several retrovirus-like proteins, including PEG10, also can form virus-like particles (VLPs), enabling the packaging of specific RNA molecules that are released from cells in extracellular vesicles (EVs) to mediate intercellular signaling.^21,22,27–34^ VLPs generated by PEG10 have even found applications in biotechnology as delivery vehicles for gene therapy and vaccines.^31,34^

Here we investigate the relationship between the retrovirus-like proteins, RTL8 and PEG10, and a key protein quality control (PQC) protein, UBQLN2. UBQLN2 is classically viewed as a ubiquitin-binding shuttle factor that interacts with and delivers polyubiquitinated client proteins to the proteasome or autophagosome for degradation.^35–37^ Recent work, however, points to a more diverse functional repertoire. For instance, UBQLN2 undergoes phase separation which allows it to modulate the fate of bound clients and may underlie its role in stress granule (SG) biology.^38–40^ UBQLN2 also interacts with chaperones, E3 ligases, and proteasome components, enabling it to regulate various aspects of PQC.^36,37,41–45^ Missense mutations in UBQLN2 directly cause familial amyotrophic lateral sclerosis (ALS) / frontotemporal dementia (FTD), further underscoring its contributions to cellular proteostasis.^46–49^

Earlier studies established that UBQLN2 facilitates proteasomal degradation of PEG10 and revealed an unexpected and novel interaction between UBQLN2 and RTL8 (Figure 1A). ^22,50,51^ In contrast to its degradative action on PEG10, UBQLN2 stabilizes RTL8, and RTL8 in turn facilitates UBQLN2 localization to the nucleus under stress conditions. Moreover, RTL8 directly interacts with PEG10, decreasing PEG10 VLP formation (Figure 1A).^33^ These results raise the intriguing possibility, examined here, of retrovirus-like proteins and UBQLN2 forming a multi-protein complex that regulates proteostasis and stress response pathways.

**Figure 1:**
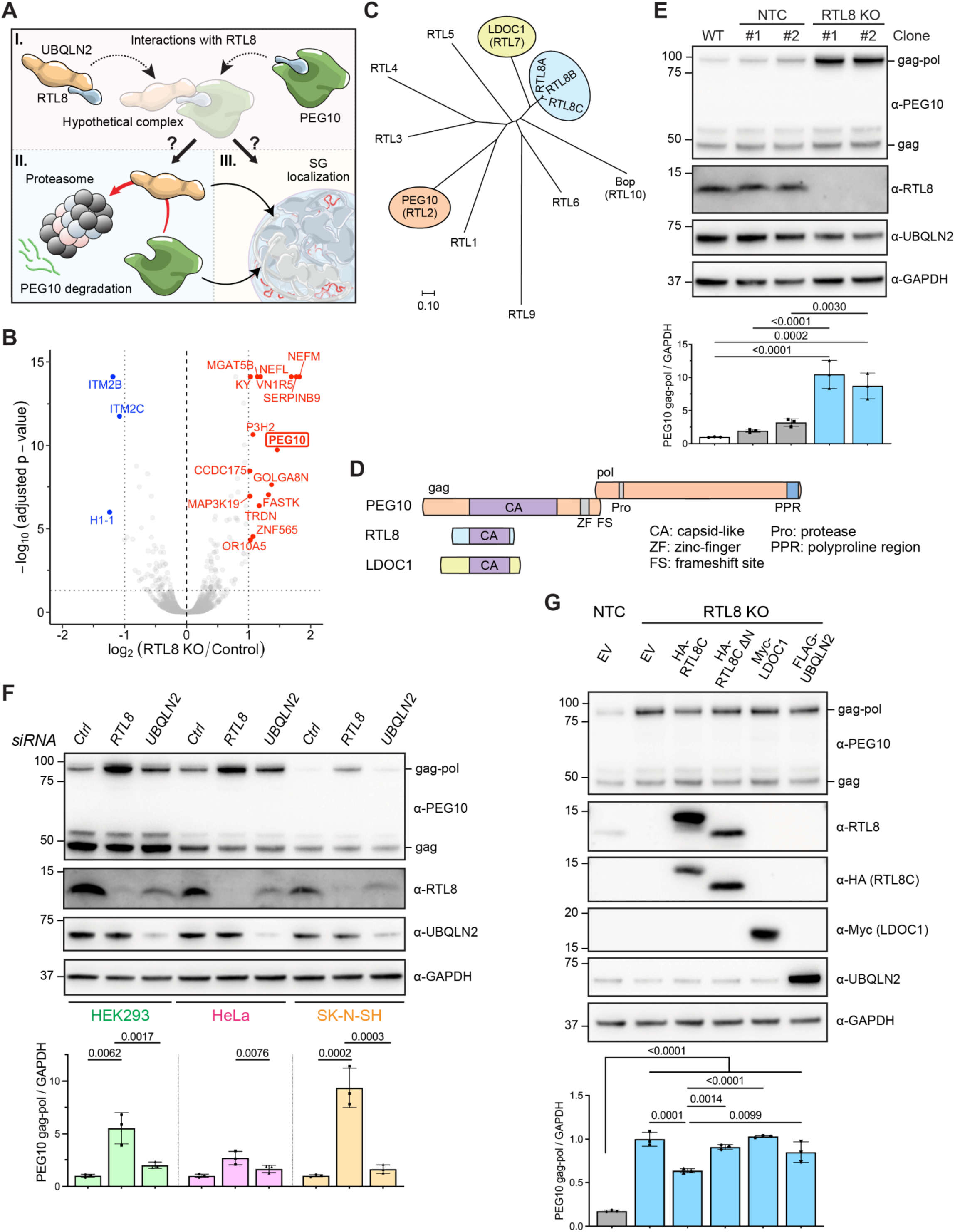
PEG10 gag-pol is regulated by the collective actions of UBQLN2 and RTL8. **A)** Schematic depicting functional relationships between UBQLN2, RTL8, and PEG10. Both UBQLN2 and PEG10 interact with RTL8 leading to a hypothetical trimolecular complex (I) that may impact UBQLN2-dependent degradation of PEG10 via the proteasome (II) and the SG localization of both proteins (III). **B)** Volcano plot of proteins significantly decreased (blue) or increased (red) in RTL8 knockout (KO) HEK293 cells compared to control cells (combination of HEK293 wild-type (WT) and non-targeting control (NTC) gRNA-treated cells). N = 3 biological replicates from 1 WT, 2 NTC and 2 RTL8 KO clonal lines. Cutoff was set at p <0.05 and absolute fold change of 2. Also, see Data S1. **C)** Phylogenetic tree of human Retrotransposon Gag-like (RTL) proteins, with branch lengths proportional to evolutionary distances. Colored ovals highlight RTL family members studied here. **D)** Domain organization of PEG10, RTL8 and LDOC1 proteins showing shared retrovirus-like features. PEG10 gag-pol is generated by ribosomal frameshifting at the frameshift site. **E)** Immunoblot of lysates from WT, NTC and RTL8 KO HEK293 cells. Relative PEG10 gag-pol levels, normalized to levels in WT cells, are shown below. Also, see Figures S1A-D. **F)** Immunoblot of lysates from various human cell lines treated with siRNA against RTL8 or UBQLN2 or control (Ctrl) siRNA. Relative PEG10 gag-pol levels, normalized to levels in control siRNA-treated cells, are shown below. Also, see Figures S1E-G. **G)** Immunoblot of lysates from NTC or RTL8 KO HeLa cells expressing HA-tagged RTL8C and RTL8C ΔN, Myc-tagged LDOC1 or FLAG-tagged UBQLN2 or empty vector. Relative PEG10 gag-pol levels, normalized to levels in RTL8 KO cells transfected with empty vector, are shown below. Also, see Figures S1H and S2. In (E-G), data represent means ± SD (N=3), analyzed with one-way ANOVA with Tukey’s multiple comparison test. GAPDH = loading control.

A feature uniting UBQLN2 and PEG10 is their ability to localize to SGs (Figure 1A).^21,38^ SGs are dynamic mRNA- and protein-containing condensates that form in response to cellular stress because of stalled translation.^52,53^ Tight regulation of SG composition, assembly, and disassembly helps maintain cellular homeostasis and ensures a rapid return to normal protein synthesis following stress relief.^54–56^ Dysregulation of SG properties likely contributes to the pathogenesis of several human diseases, including cancer and neurodegenerative disorders like ALS.^53,54,57–59^ Defining how the interplay between retrovirus-like proteins and UBQLN2 affects stress-induced condensate formation and function may provide insight into the pathological processes underlying these diseases.

Here we establish that PEG10 and RTL8 functionally converge on UBQLN2 to modulate pathways of proteostasis and stress response. RTL8 enables UBQLN2-mediated turnover of PEG10. Both proteins are required to recruit UBQLN2 to SGs, with RTL8 serving as an essential bridge connecting UBQLN2 to PEG10. Changes in the levels of any of the three proteins, especially PEG10, result in significant changes to SG dynamics and overall composition. Our further discovery of PEG10-derived VLPs within SGs underscores the heterogeneity and structural complexity of SGs, suggesting previously unrecognized roles for retrovirus-like proteins in the organization and function of SGs during stress responses.

## MATERIALS AND METHODS

### Key resources table

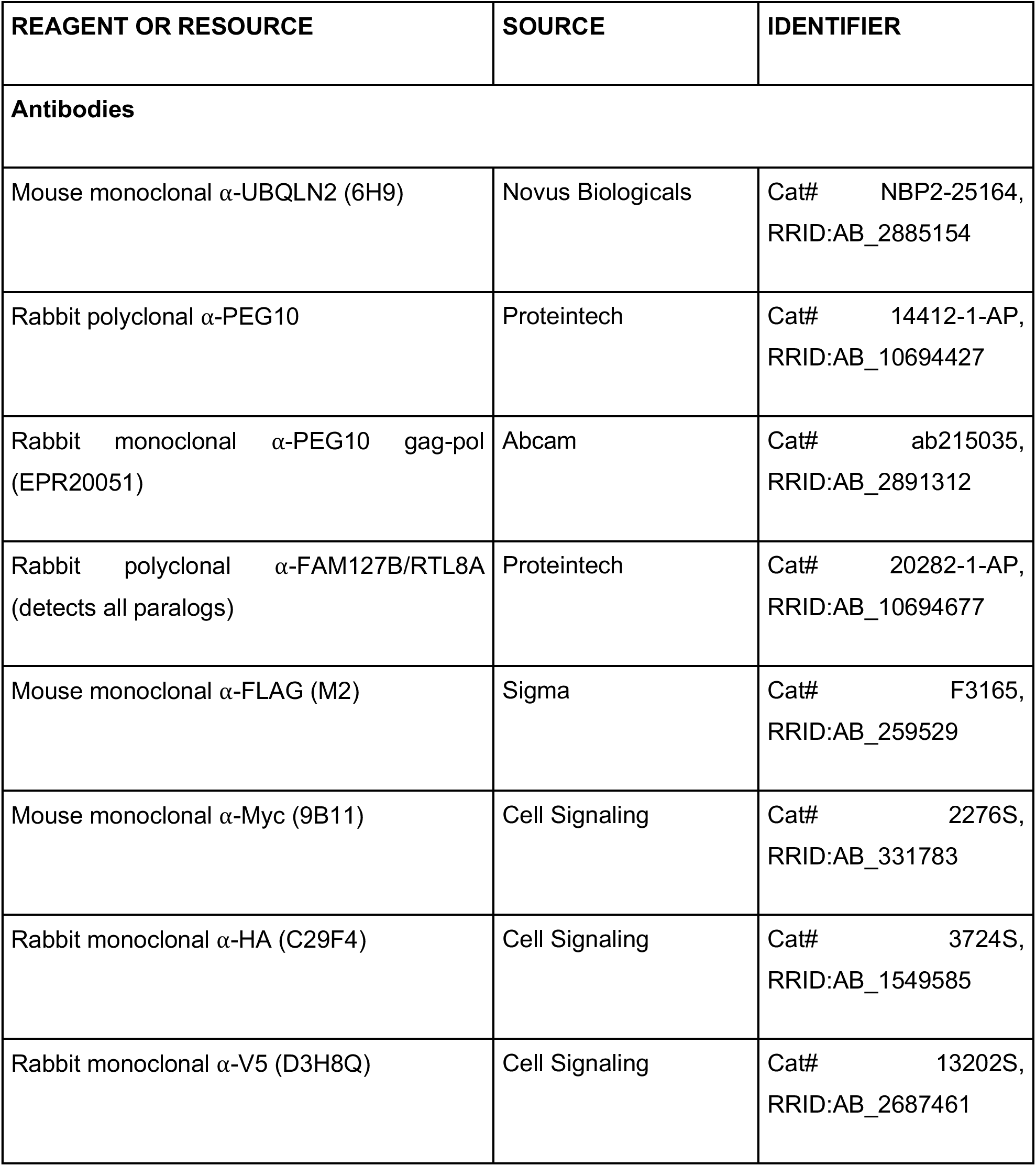
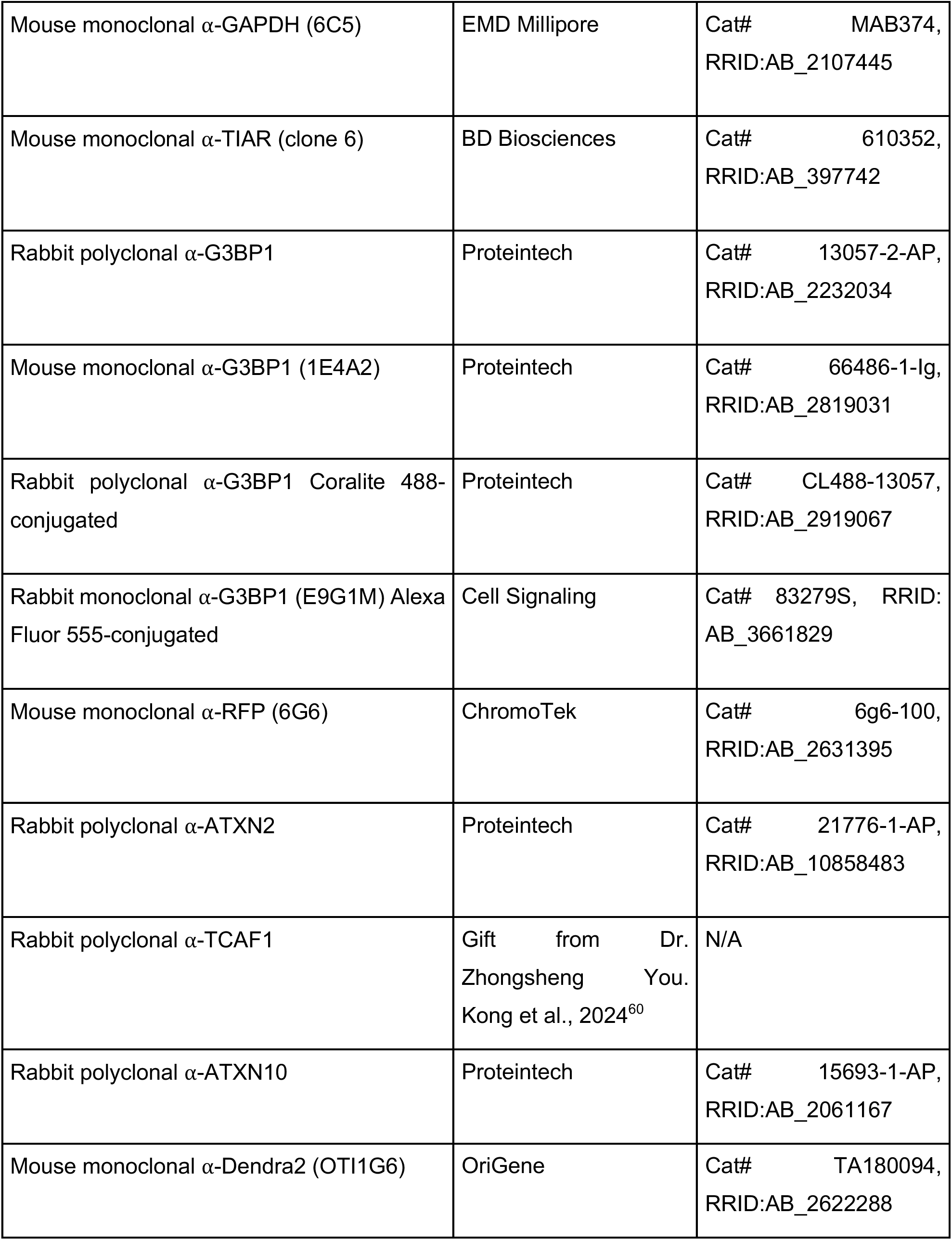
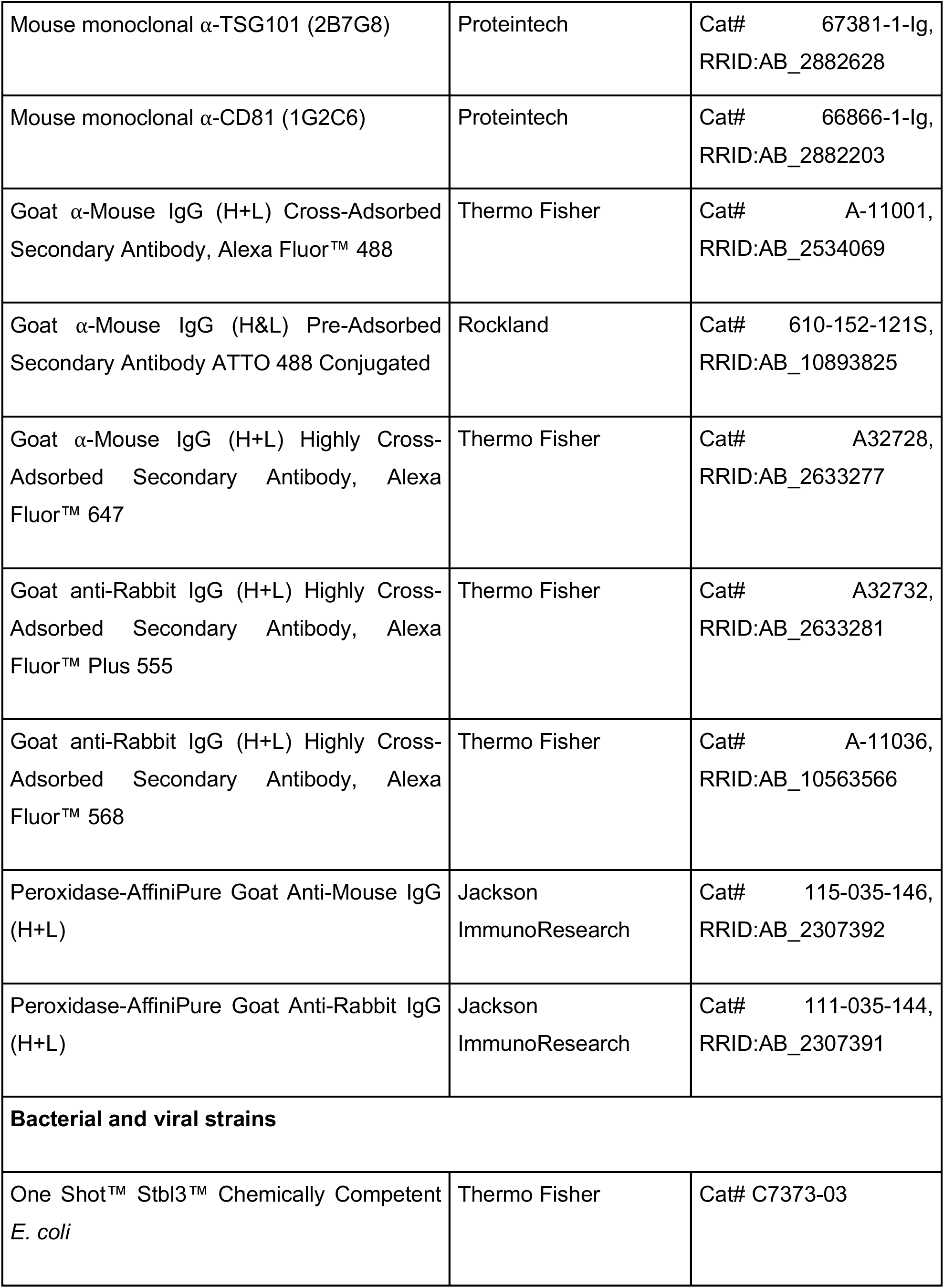
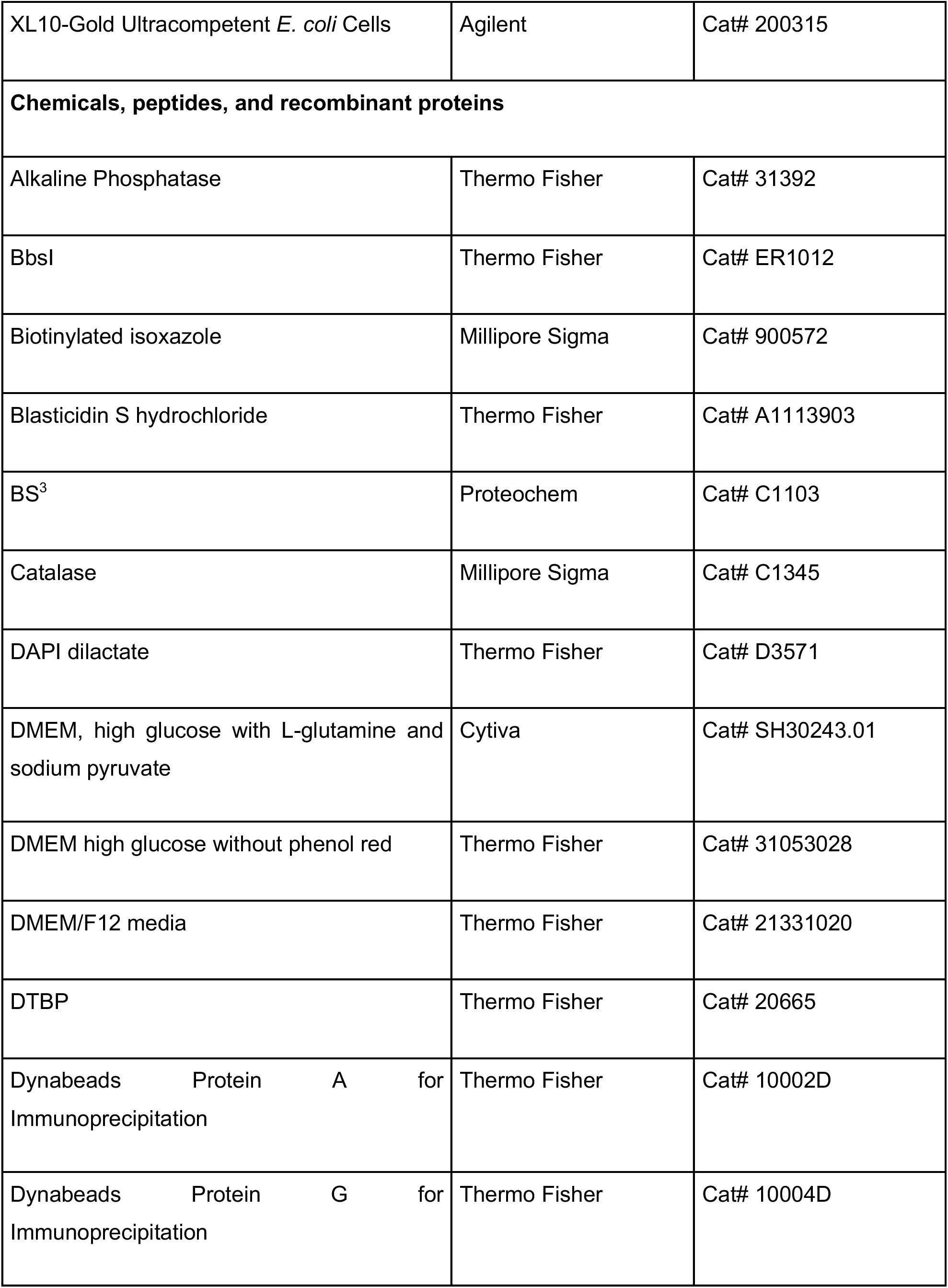
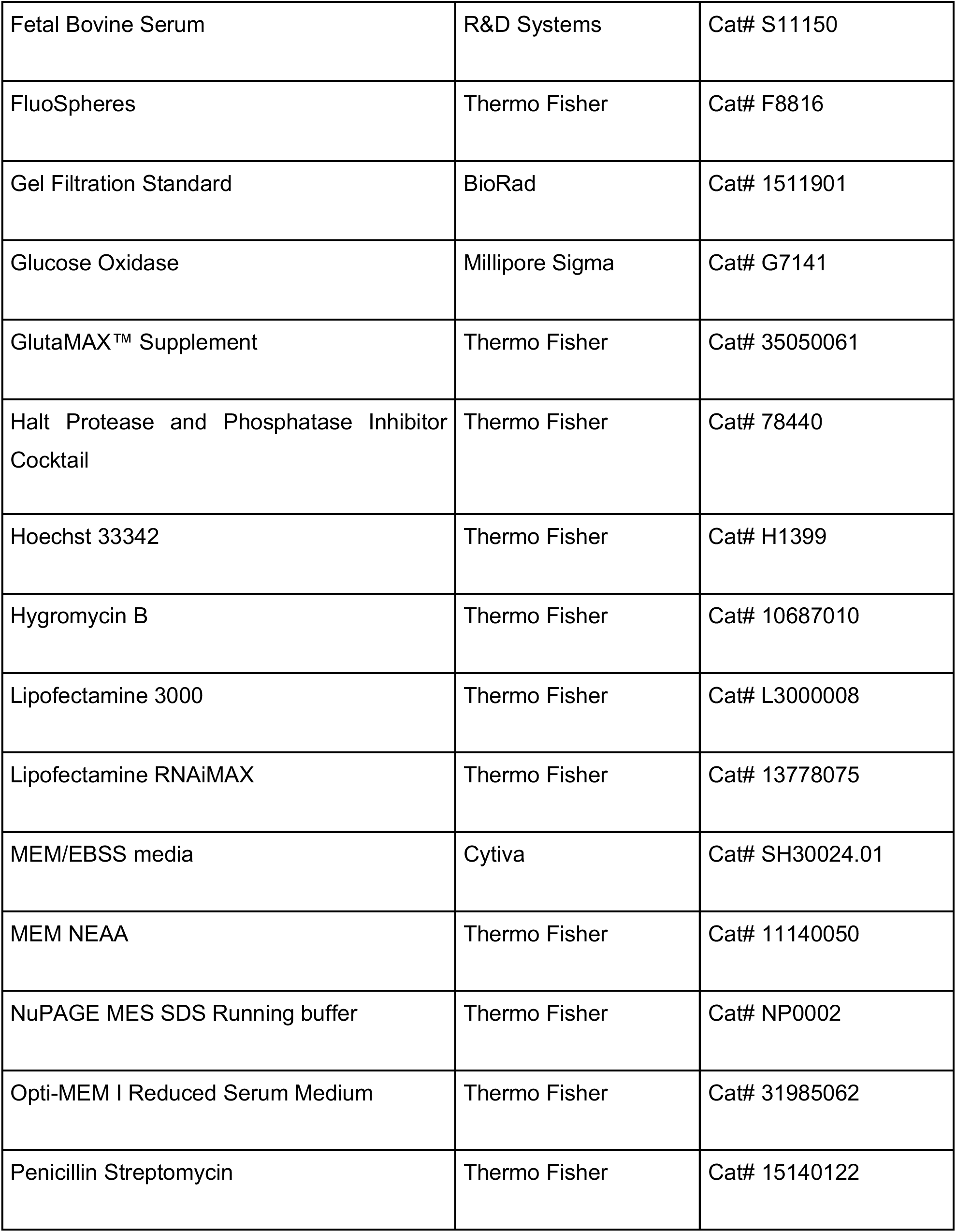
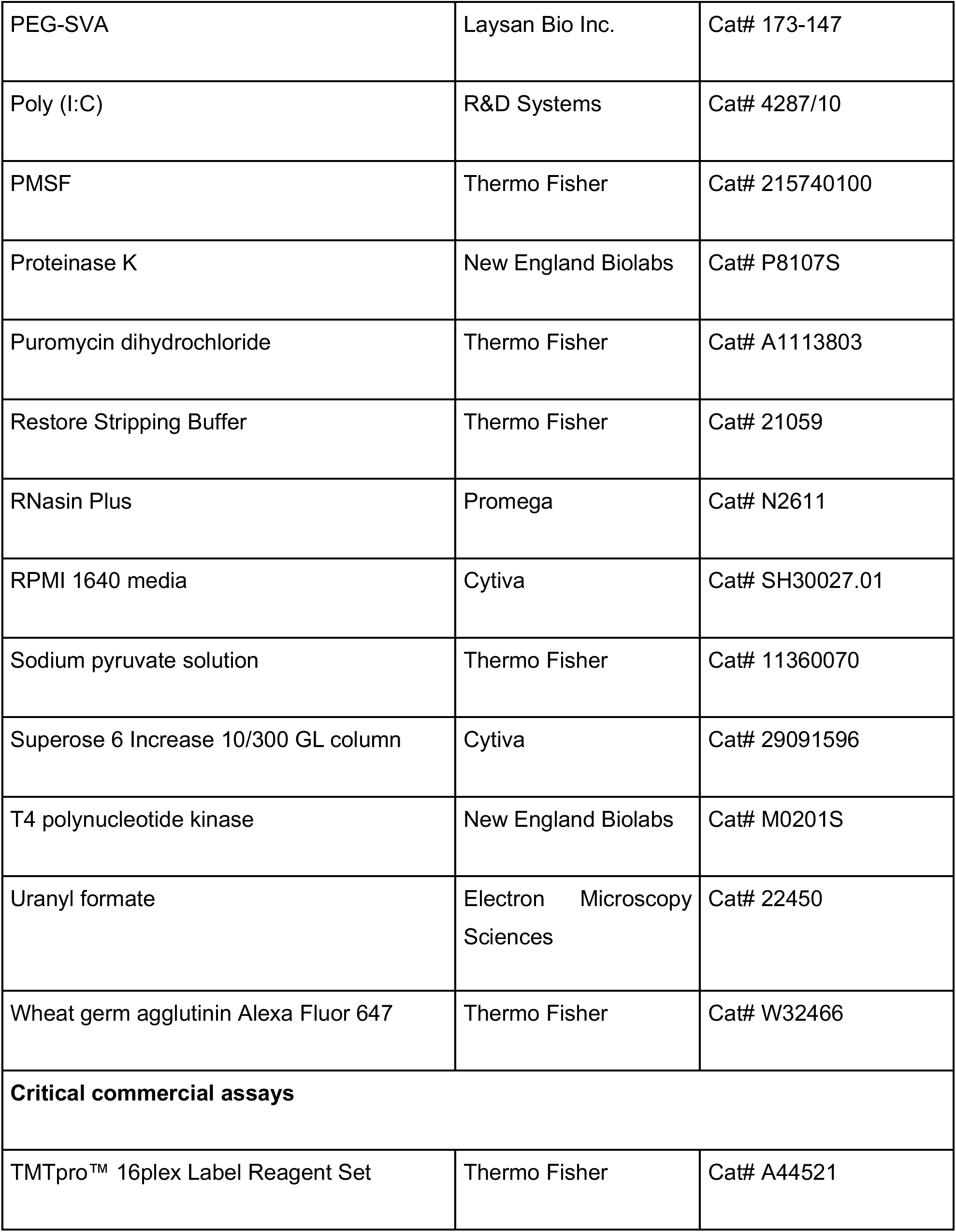
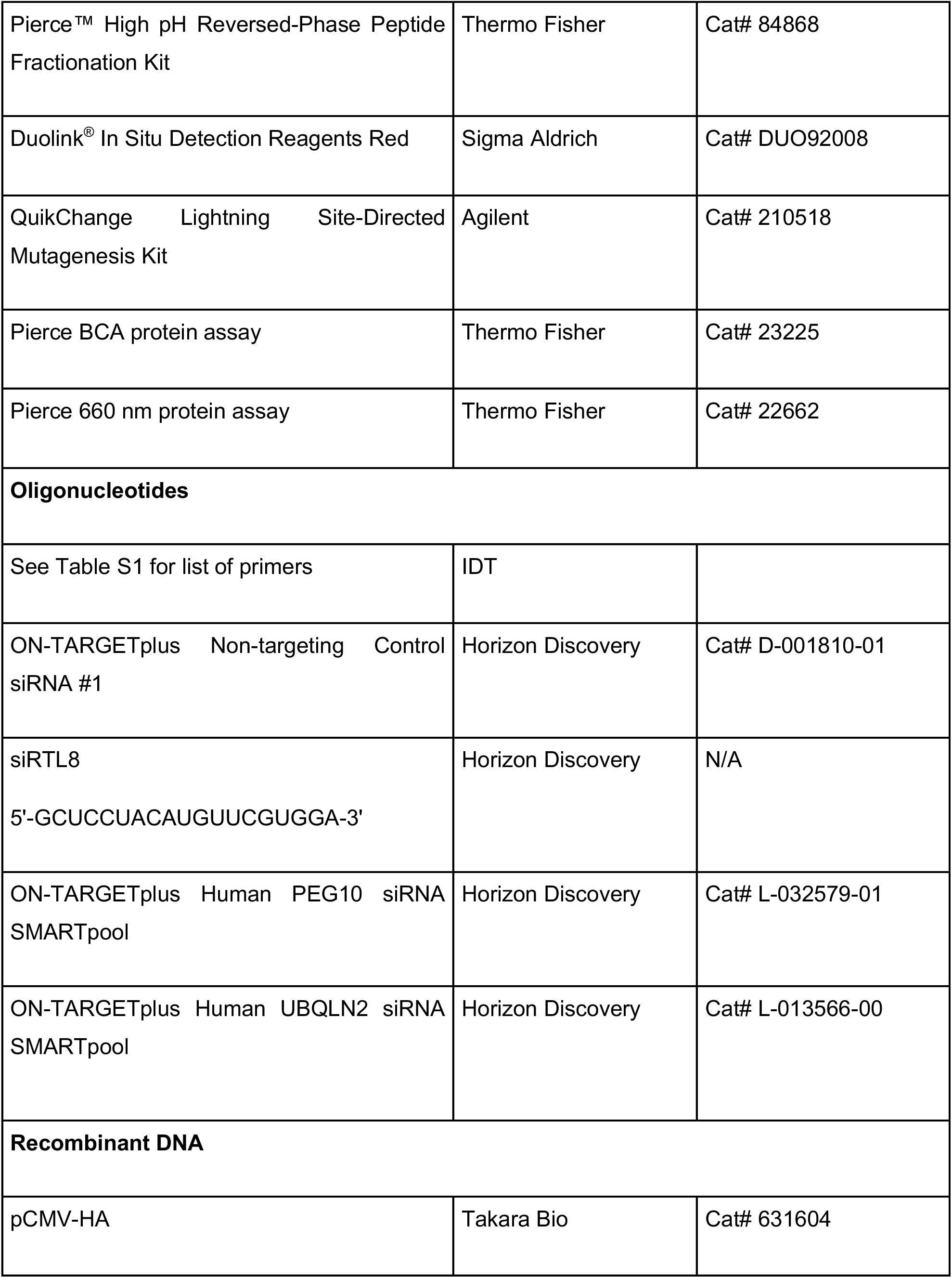
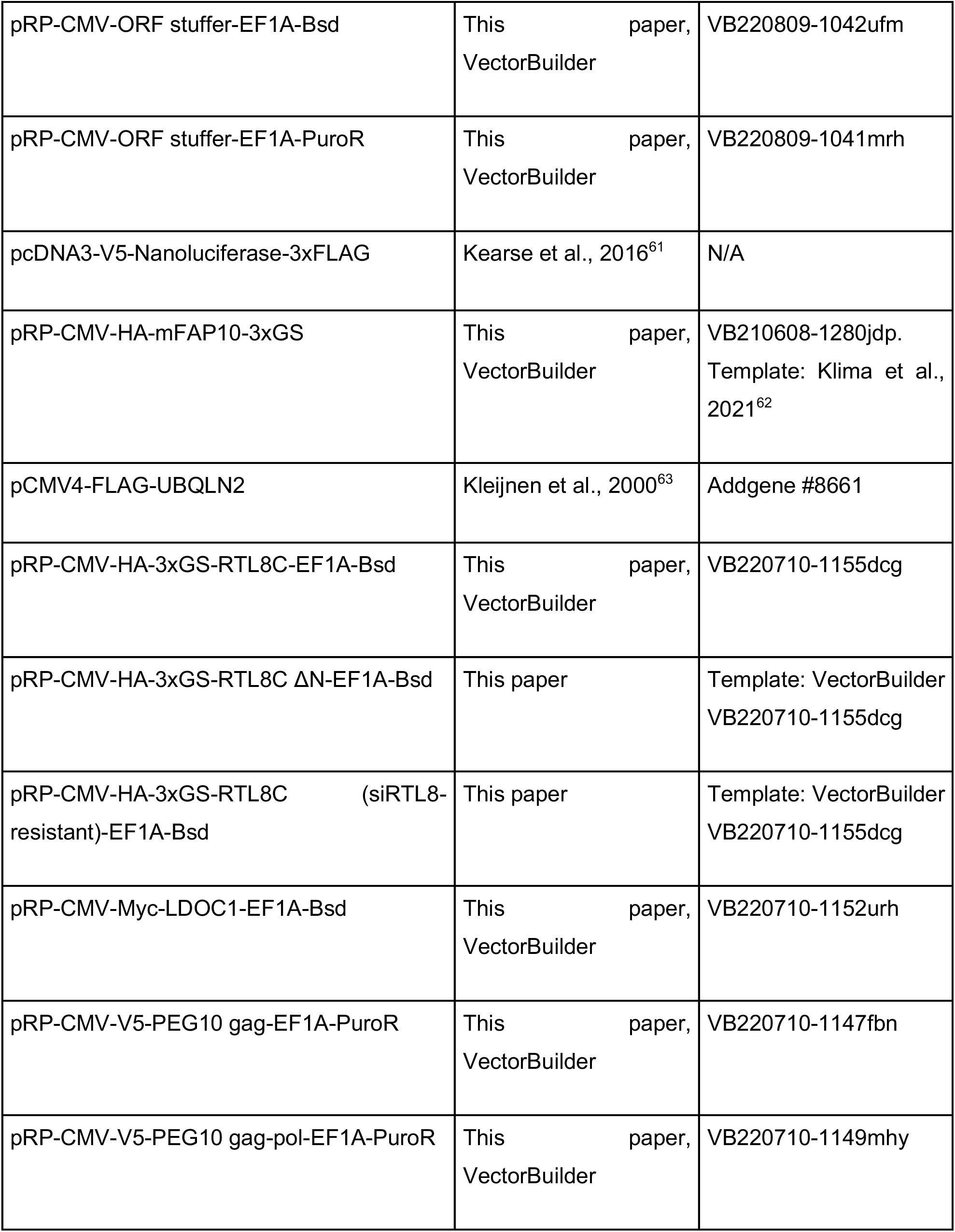
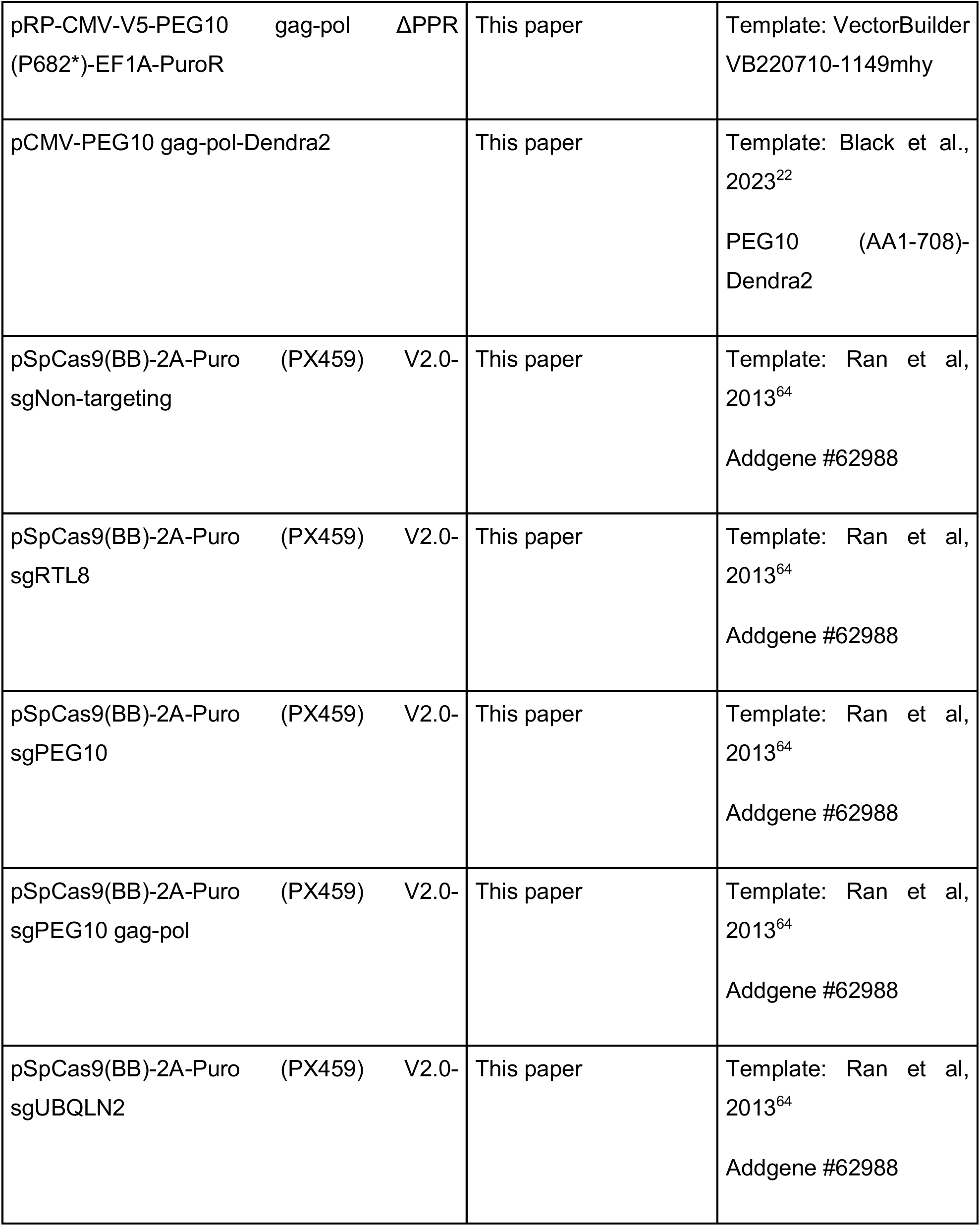
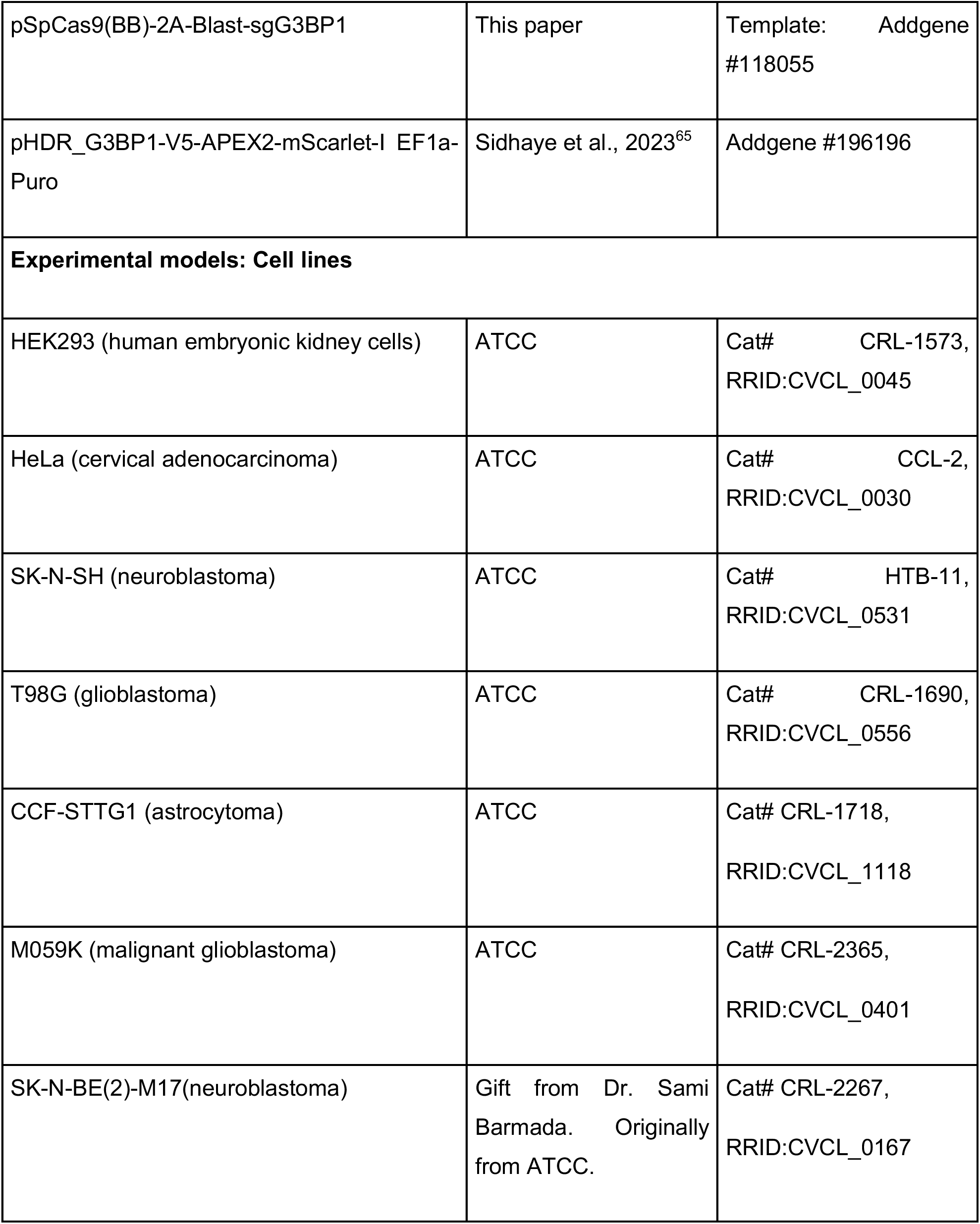
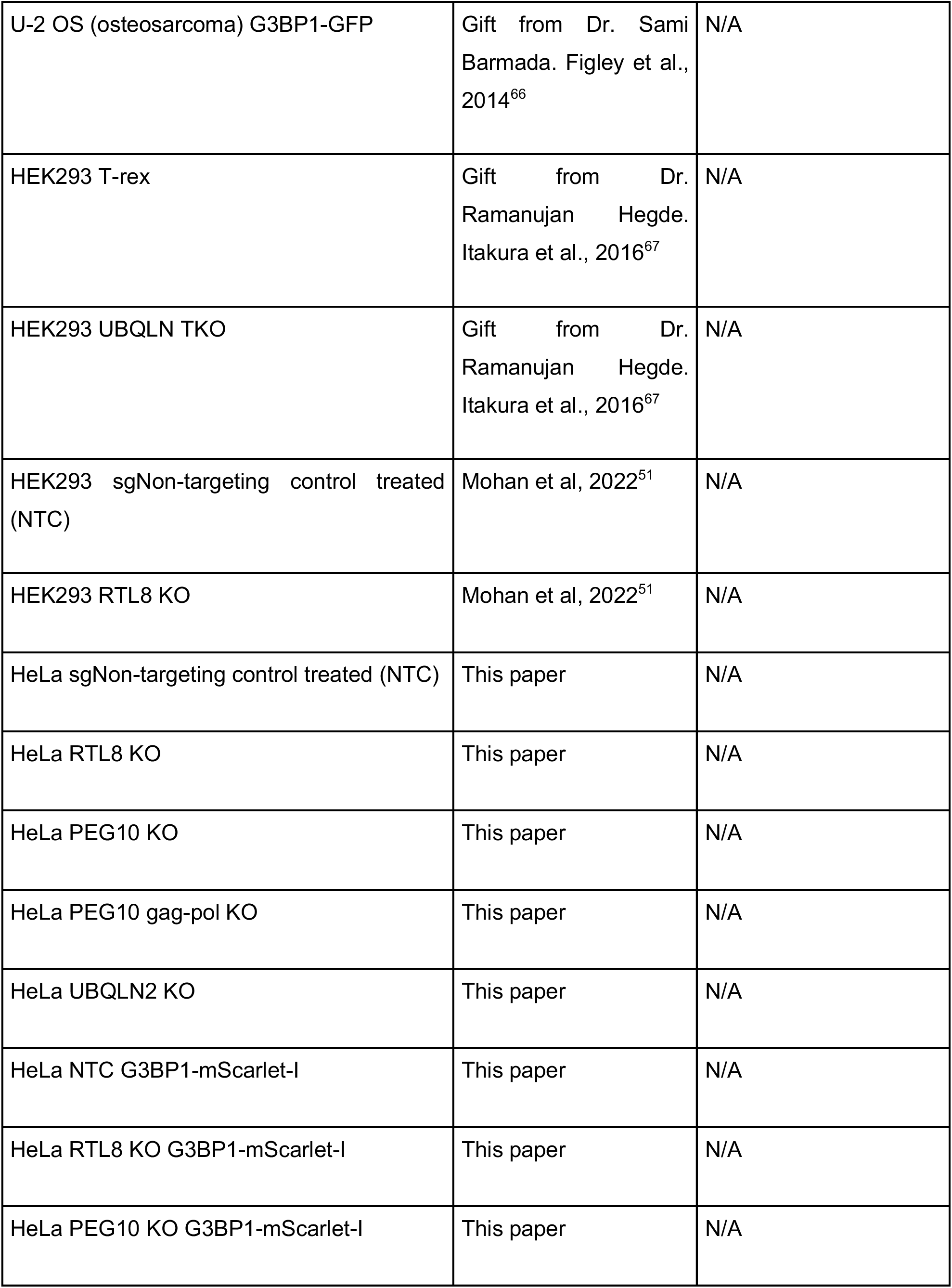
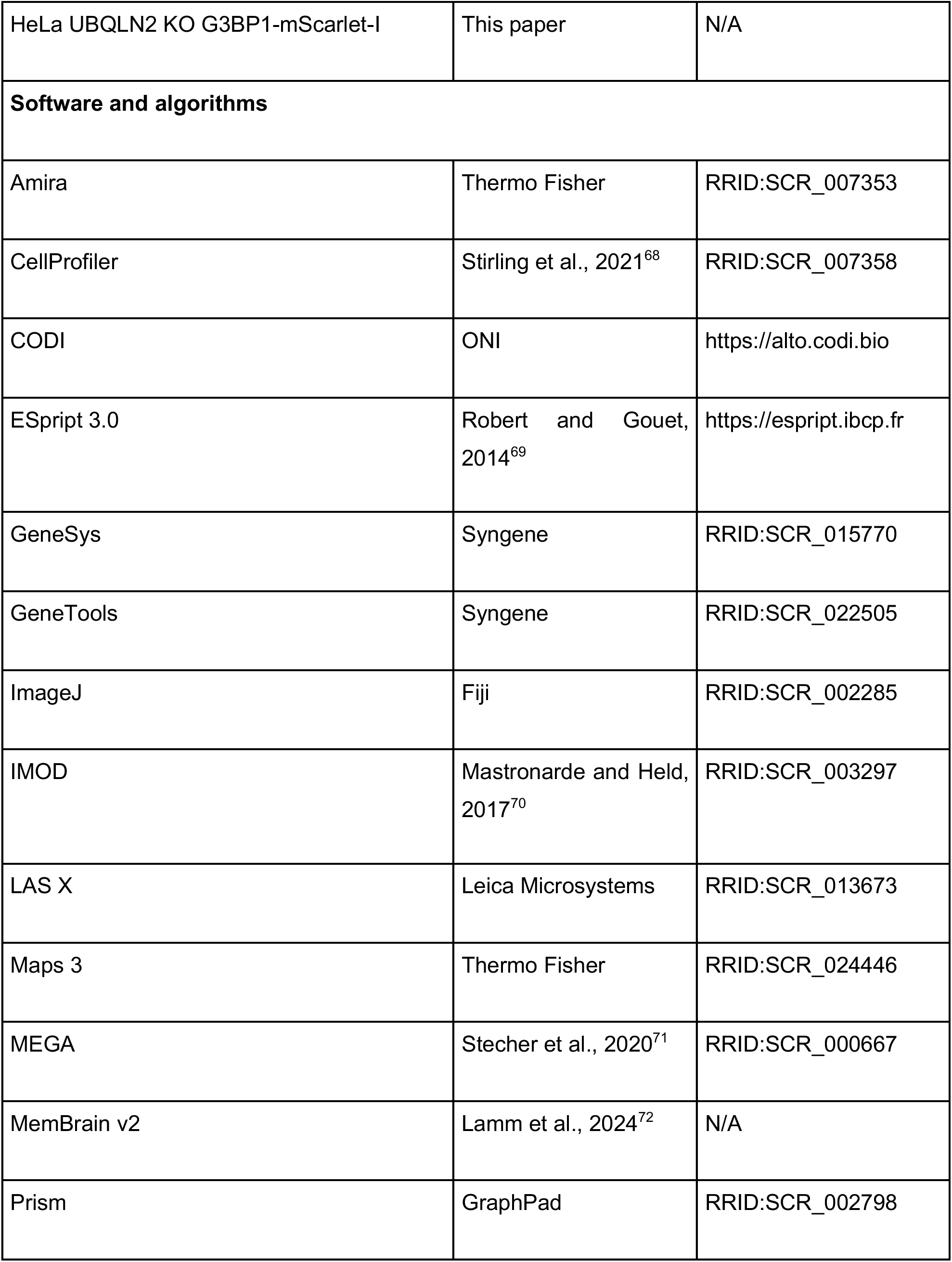
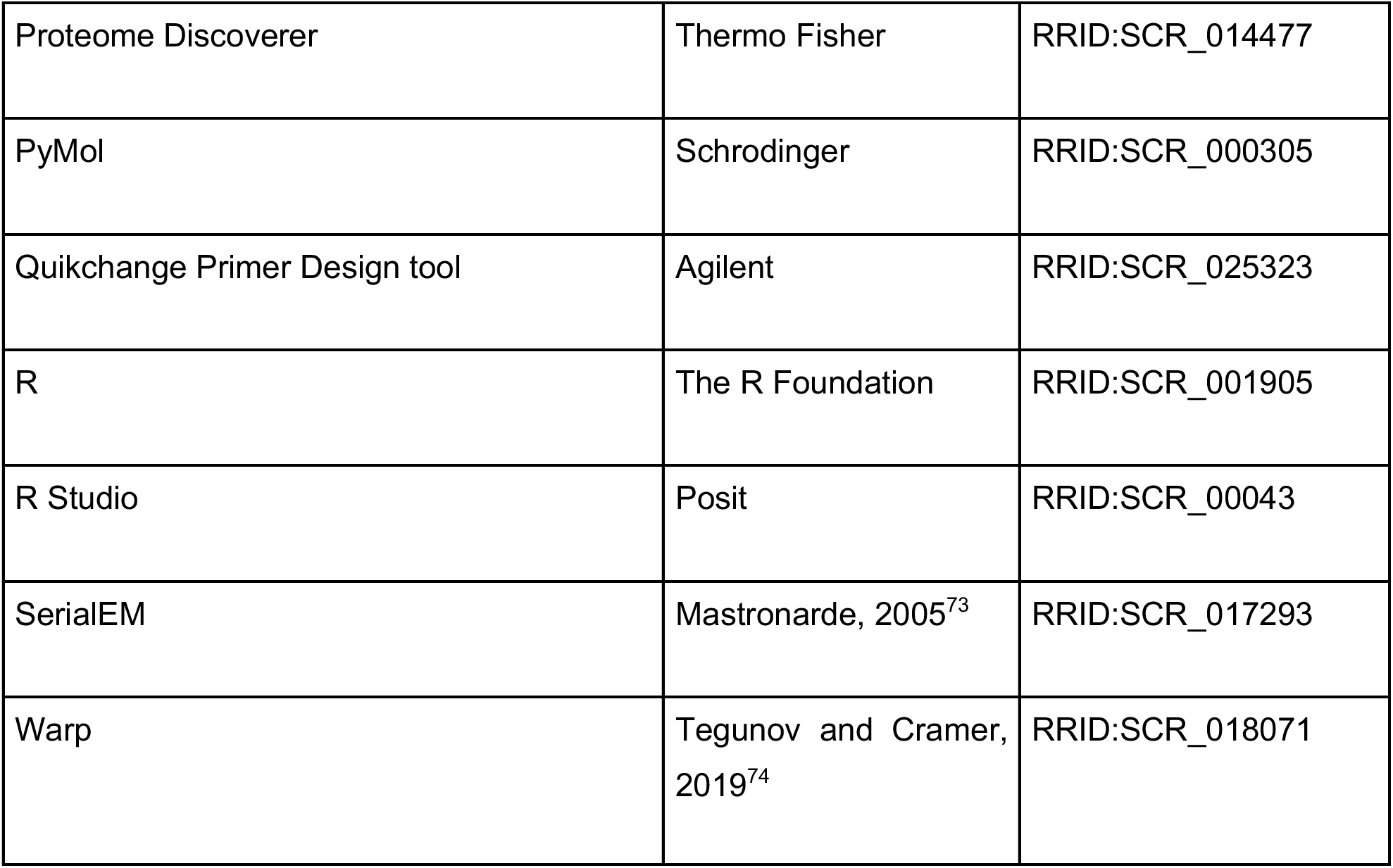

### Cloning and site-directed mutagenesis (SDM)

Plasmids for mammalian gene expression were codon-optimized and synthesized by VectorBuilder. Plasmids were verified by Sanger sequencing prior to use.

Non-targeting guide or guide sequences targeting RTL8, PEG10, PEG10 gag-pol, or UBQLN2 were cloned into pSpCas9(BB)-2A-Puro (PX459, Addgene, 62988) and guide sequence targeting G3BP1 was cloned into cloned into pSpCas9(BB)-2A-Blast (Addgene, 118055). Guide sequences were cloned following a previously published protocol (Ran et al., 2013). Briefly, PX459 was digested using BbsI (Thermo Fisher), dephosphorylated using alkaline phosphatase (Thermo Fisher) and gel purified. DNA oligos containing the guide sequences (IDT) were annealed and phosphorylated using T4 polynucleotide kinase (NEB). Purified digested plasmids and annealed oligos were ligated and transformed into Stbl3 competent cells (Thermo Fisher). Clones were picked and Sanger sequenced using U6 promoter FP.

SDM was performed using Quikchange Lightning kit (Agilent) as per manufacturer’s instructions. Primers for SDM were designed using the Quikchange Primer Design tool (Agilent). RTL8C ΔN was generated by deleting the nucleotides between residues Asp2 and Asn25. siRTL8 resistant HA-RTL8C was designed using the Synonymous Mutation Generator tool.^75^ To generate a plasmid expressing only PEG10 gag-pol-Dendra2, an adenine was inserted at position 1568 in the PEG10 gag-pol ORF to force the frameshift and Glu11 in the CFP ORF was mutated to a stop codon. Overlap extension PCR was done using 50 ng of template and 125 ng of each primer on a thermocycler with the following conditions: 95°C for 2 min, 18 cycles of 95°C for 20 s, 60°C for 10 s, 68°C for 3-4 min and final extension at 68°C for 5 min. DpnI was added to the mixture and incubated for 10 minutes at 37°C. The mixture was transformed into XL10 Gold ultracompetent cells (Agilent). Clones were picked and Sanger sequenced using PEG10 sequencing primers and IRES seq F1 for PEG10 constructs and CMVf for RTL8 constructs.

### Mammalian cell culture

HEK293 (ATCC) and HeLa (ATCC) cell lines were maintained in complete DMEM (DMEM containing L-glutamine and sodium pyruvate (Cytiva Life Sciences) supplemented with 10% FBS (R&D Systems) and 100 U/mL Penicillin-Streptomycin (Thermo Fisher)). For live cell imaging studies, HeLa cells were grown in complete DMEM without phenol red. SK-N-SH and T98G (both ATCC) cell lines were maintained in MEM/EBSS (Cytiva Life Sciences) supplemented with 1x GlutaMAX (Thermo Fisher), 1 mM sodium pyruvate (Thermo Fisher), 0.1 mM NEAA (Thermo Fisher), 10% FBS and 100 U/mL Penicillin-Streptomycin. CCF-STTG1 cell line (ATCC) was maintained in RPMI-1640 (Cytiva Life Sciences) supplemented with 10% FBS and 100 U/mL Penicillin-Streptomycin. M059K and BE(2)-M17 (both ATCC) cell lines were maintained in DMEM/F12 (Thermo Fisher) supplemented with 10% FBS and 100 U/mL Penicillin-Streptomycin. U-2 OS cells stably expressing G3BP1-GFP were maintained in complete DMEM. HEK293 T-rex control and UBQLN1, UBQLN2 and UBQLN4 knockout (TKO) cell lines were maintained in complete DMEM supplemented with 10 μg/mL Blasticidin S (Thermo Fisher), and 100 μg/mL Hygromycin (Thermo Fisher). All lines were maintained at 37°C and 5% CO_2_.

### Stress conditions

Stress experiments were carried out as follows. Cells were heat shocked by placing them in an incubator preset to 43°C for 1 h. Oxidative stress was performed by adding complete DMEM containing 0.5mM sodium arsenite to cells for 45 minutes. Hyperosmotic stress was induced by adding complete DMEM containing 0.2 M sodium chloride to cells for 45 minutes. dsRNA stress was initiated by transfecting cells with poly (I:C) (R&D Systems) at a final concentration of 0.5 ng/µL using Lipofectamine 3000 (Thermo Fisher) for 6 h as outlined below.

### Transfection

All transfections were performed 24h after cells were plated. Plasmids and poly (I:C) were transfected using Lipofectamine 3000, 24 h after cells were plated. Plasmids were diluted in Opti-MEM I Reduced Serum Medium (Thermo Fisher) along with P3000 at a ratio of 2 µL of P3000 per µg of DNA. For microscopy experiments, P3000 was omitted. Separately, Lipofectamine 3000 was diluted in Opti-MEM at a ratio of 1.5-2 µL of Lipofectamine 3000 per µg of DNA. The Lipofectamine 3000 mixture was combined with the plasmid-P3000 mixture, incubated for 15 minutes at room temperature, and added to cells. Media containing the transfection mixture was replaced with fresh media 6 h after transfection. Cells were used for downstream analyses 24-48 h post transfection. For poly (I:C) transfections, P3000 was omitted and 3 µL of Lipofectamine 3000 was used per µg of poly (I:C) in Opti-MEM. For generation of stable lines expressing an empty vector or HA-RTL8C, cells were selected with media containing 10 µg/mL Blasticidin for 3 days, following which clones were isolated via limiting dilution (1 cell/well) in 96-well plates, and RTL8 levels were verified by immunoblotting.

For siRNA transfections, Lipofectamine RNAimax (Thermo Fisher) was used at a ratio of 0.3 µL of Lipofectamine RNAimax per picomole of siRNA. Lipofectamine RNAimax and siRNA were diluted separately in OptiMEM. The mixtures were combined at a 1:1 ratio by volume, incubated for 5 minutes at room temperature, and added to cells such that the final concentration of siRNA per well was 10 nM. Cells were used for downstream analyses 48 h post transfection.

For siRNA and plasmid co-transfections, P3000 was omitted and Lipofectamine 3000 was diluted in Opti-MEM at a ratio of 1.5 µL of Lipofectamine 3000 per µg of nucleic acid (the amounts of plasmid and siRNA were 1 and 0.15 µg, respectively). The Lipofectamine 3000 mixture was combined with the siRNA-nucleic acid mixture, incubated for 15 minutes at room temperature, and added to cells. Cells were used for downstream analyses 48 hours post transfection.

### Generation of CRISPR knockout and knock-in lines

HEK293 non-targeting control (NTC) and RTL8 KO lines were generated previously (Mohan et al, 2022^51^). For gene knockouts in HeLa cells, the generated plasmids were then transfected into HeLa cells. 24 h after transfection, cells were selected with 0.4 μg/mL puromycin for 5-7 days. Clones were isolated via limiting dilution (1 cell/well) in 96-well plates, and knockout was verified by immunoblotting for RTL8, PEG10, or UBQLN2. The guide RNA sequences were as follows:

sgNon-targeting: 5’-GATACGTCGGTACCGGACCG-3’

sgRTL8: 5’-GTTCTCGTCCACGAACATGT-3’

sgPEG10 gag-pol: 5’-GATCATGGCTCGGACGAACA-3’

sgPEG10: 5’-GCTGATTGACCAGTACCACG-3’

sgUBQLN2: 5’-ACGCAGCCTAGCAATGCCGC-3’

sgG3BP1: 5’-TCCATGAAGATTCACTGCCG-3’

For CRISPR knock-in of a V5-APEX2-mScarlet-I tag into the endogenous G3BP1 locus, the generated Cas9-sgG3BP1 plasmid (template Addgene #118055) was co-transfected with the HDR template pHDR_G3BP1-mScarlet (Addgene #196196) at a 1:1 ratio into the various HeLa KO lines. 48 h after transfection, cells were treated with 0.4-0.5 μg/mL puromycin (Thermo Fisher) for 3-4 days to select for successful integration of the insert, which confers puromycin resistance. Surviving fluorescent colonies were then expanded and knock-in was confirmed by immunoblotting.

### Preparation of cell lysates, immunoblotting and quantification

Unless otherwise mentioned, cell lysates were prepared as follows. Cells were harvested by trypsinization or by scraping in ice-cold 1x phosphate-buffered saline (PBS), washed in ice-cold 1x PBS and pelleted by centrifuging at 2200xg for 1 minute at 4°C. Cell pellets were resuspended in Pierce RIPA buffer (Thermo Fisher) supplemented with Halt™ protease and phosphatase inhibitors and 5 mM EDTA (Thermo Fisher), sonicated on ice for 10 seconds and centrifuged at 16,000xg for 20 minutes at 4°C. The supernatants were collected, and Pierce BCA or 660 nm assay (Thermo Fisher) was performed to estimate protein concentration. Equal amounts of protein from each sample was combined with 6x Laemmli buffer containing 600 mM DTT to a final concentration of 1x and boiled for 10 minutes.

Protein samples (lysates and IP eluates) were resolved by gel electrophoresis on 4-12% NuPAGE Novex Bis-Tris gels (Thermo Fisher) using NuPAGE MES SDS running buffer (Thermo Fisher) at 140V. Protein transfer was performed at 110V for 1h onto 0.2 μm nitrocellulose membrane (Biorad). Membranes were stained in 0.2% Ponceau S (Thermo Fisher) in 5% acetic acid (Thermo Fisher) to ensure uniform loading. They were washed in 1x Tris-buffered saline containing 0.1% Tween-20 (Thermo Fisher) (TBST) and subsequently blocked in blocking buffer comprising of 5% non-fat dry milk (DotScientific), 0.05% bovine serum albumin (BSA, Thermo Fisher), and 0.01% thimerosal (Thermo Fisher) in 1x TBST for 1 h at RT. Membranes were then incubated overnight at 4°C with primary antibodies diluted in blocking buffer with gentle agitation, then washed thrice with 1x TBST and incubated with secondary antibodies for 1 h at room temperature. After washing with 1x TBST, blots were developed using EcoBright Nano or Femto HRP (Innovative Solutions) on a G Box Mini 6 imager (Syngene). Blots were stripped with Restore Stripping Buffer (Thermo Fisher), washed with TBST and reprobed.

GeneTools (Syngene) was used for blot quantification. Unless otherwise mentioned, band intensities were normalized to their respective GAPDH lanes.

### Immunocytochemistry, dSTORM and proximity ligation assay (PLA)

Cells were seeded onto poly-D-lysine pre-coated coverslips (Neuvitro Corporation) at least 24 h prior. After stressing, coverslips were washed once in PBS and fixed with 4% paraformaldehyde (Electron Microscopy Sciences) in PBS for 10 minutes at RT. Unreacted paraformaldehyde was quenched by incubating in 50 mM glycine in PBS for 5 minutes. Coverslips were washed thrice in PBS, blocked and permeabilized in blocking buffer consisting of 5% normal goat serum (Vector Laboratories), 0.05% BSA and 0.01% Triton X-100 (Thermo Fisher) in PBS, and then incubated with primary antibodies diluted in blocking buffer overnight at 4°C with gentle agitation. Coverslips were then washed in PBS containing 0.1% Tween-20 (PBST) and incubated with secondary antibodies in blocking buffer containing 1 µg/mL DAPI dilactate (Thermo Fisher), followed by washes in PBST. They were mounted using Prolong Glass (Thermo Fisher) and Z-stacks with 0.12 µm spacing were obtained on the Stellaris 5 confocal microscope (Leica Microsystems) with a 63x or 100x oil objective. Maximum intensity projections (MIPs) of Z-stacks were generated on LAS X (Leica Microsystems) or ImageJ. Images were analyzed using custom CellProfiler pipelines that have been deposited. Briefly, nuclei and cytoplasm were identified by DAPI and diffuse G3BP1 staining, respectively; SGs were identified by intense G3BP1 puncta and used for masks. Intensities, areas and Mander’s overlap coefficients were calculated using CellProfiler modules. Intensities of proteins in SGs were baseline corrected to their respective cytoplasmic intensities.

For dSTORM imaging, cells were seeded onto Lab Tek-II chamber slides (Thermo Fisher) 24h prior to staining. Cells were stressed with 0.5 mM sodium arsenite, fixed and stained as mentioned above and stored in PBS. Prior to imaging, slides were washed once with buffer containing 100 mM Tris HCl pH 8 and 25 mM NaCl and imaged in GLOX buffer (100 mM Tris HCl pH 8, 25 mM NaCl, 10% glucose, 142 mM β-mercaptoethanol, 80 μg/mL catalase, and 0.5 mg/mL glucose oxidase) on the NanoImager. Images were analyzed on CODI (ONI). Jaccard index between channels was calculated by dividing the number of clusters that were positive for both channels by the total number of clusters positive for either channel.

For PLA experiments, cells were seeded and fixed as described above. Cell membrane staining was done by incubating coverslips in 5 µg/mL wheat germ agglutinin-Alexa Fluor 647 (Thermo Fisher) in HBSS for 10 minutes at RT. Coverslips were then washed thrice in PBS and PLA was carried out using Duolink PLA Red kit (Sigma) as per manufacturer’s protocol with the following modifications: 1) primary antibodies were diluted 1:2000 in Duolink blocking solution and incubated for 2 h at RT; 2) secondary probes were diluted 1:10 in Duolink blocking solution; and 3) Amplification was carried out for 1 h at 37°C. Coverslips were mounted and imaged as described above. PLA foci per cell and with respect to area were calculated using custom CellProfiler pipelines. Cell nuclei were identified by DAPI staining and cytoplasm by WGA.

### Live cell imaging

G3BP1-mScarlet tagged HeLa cell lines were seeded onto chambered coverslips (Ibidi) and imaged after 48 hrs. Nuclei were labeled with Hoechst 33342 (Thermo Fisher) at a final concentration of 1 ug/mL in media for 20 min, followed by 2 media washes. Confocal imaging was performed at 20x magnification with the Leica Mica microscope at 37 °C, 63% humidity, and 5% CO2. For SG assembly, cells were stressed with 0.5 mM sodium arsenite for 20-24 min with timelapse imaging every 2 min. For SG disassembly, stressed cells were washed with fresh media once, then imaged every 10 min for 2.5 hrs. Images were analyzed using CellProfiler. Nuclei and cytoplasm were identified by Hoechst and diffuse G3BP1 staining, respectively. SGs were identified by intense G3BP1 puncta. Average number of SGs per cell per field were calculated at each time point, normalized to the peak SG count per cell for the field over the course of imaging, and graphed. SG size and intensity were quantified at 20 minutes post arsenite treatment.

### Co-immunoprecipitation (co-IP) and SG IP

Cells were harvested 48h after seeding or transfection from 15 cm plates. 5 µg of antibodies were irreversibly crosslinked to Dynabeads Protein A or Protein G (Thermo Fisher) using 5 mM BS^3^ (ProteoChem) and excess crosslinker was quenched using 1 M Tris pH 7.6. Crosslinked beads were washed in PBST and stored at 4°C until use. Cells were trypsinized and washed in ice cold PBS. Crosslinking IP was performed as described in ^76^. Briefly, cells were incubated in 4 mM DTBP (Thermo Fisher) for 1h on ice and excess DTBP was quenched using 1 M Tris pH 7.6. Cells were then lysed in IP lysis buffer (50 mM Tris pH 7.6, 150 mM NaCl, 1% Triton X-100, 5% glycerol, 5 mM EDTA with protease and phosphatase inhibitors (Thermo)) for 30 minutes at 4°C with rotation. Lysates were spun at 16,000xg for 20 minutes at 4°C. Supernatants were collected and precleared using control IgG-conjugated beads. Protein quantitation was done on pre-cleared supernatants using a Pierce 660 nm assay (Thermo Fisher); 1 mg and 10 µg of protein were used for IP and input, respectively. Antibody-conjugated beads were added to IP samples which were rotated for 2h at 4°C. Samples were then washed thrice with IP lysis buffer, resuspended in 1x Laemmli buffer containing 100 mM DTT and boiled at 95°C for 10 minutes to elute bound proteins.

SG cores were isolated from HeLa cells as described previously.^77,78^ Briefly, cells were plated onto 500 cm^2^ dishes (Corning) and stressed with sodium arsenite 48h later. Cells were washed with warm media, scraped and pelleted by centrifugation, then snap frozen in liquid nitrogen and thawed on ice prior to lysis. All subsequent steps were carried out at 4°C. Cells were lysed in SG IP buffer (50 mM Tris pH 7.6, 50 mM NaCl, 5 mM MgCl_2_, 0.5% NP40, 0.5 mM DTT and 50 µg/mL heparin sodium (Sigma) supplemented with protease and phosphatase inhibitors and 100 U/mL RNaseIN Plus (Promega)) by passing through a 25G needle 7 times and incubated for 15 minutes with rotation, then centrifuged at 1000xg for 5 minutes. The supernatants were collected and spun at 18,000xg for 20 minutes. The pellets were retained and resuspended in SG IP buffer, then spun at 850xg for 2 minutes after which the supernatant (SG core-enriched fraction) was collected. SG core-enriched fractions were precleared using rabbit IgG-conjugated Dynabeads. Approximately 200 µg of the precleared SG core-enriched fractions as determined by a Pierce 660 nm assay (Thermo Fisher) were used for IP. SG cores were immunoprecipitated by incubating with rabbit α-G3BP1-conjugated Dynabeads for 4h with rotation. Beads were then washed as follows: 1) twice for 5 minutes with rotation in Wash buffer 1 (20 mM Tris pH 8, 200 mM NaCl with protease and phosphatase inhibitors and 100 U/mL RNaseIN Plus); 2) 5 minutes with rotation in Wash buffer 2 (20 mM Tris pH 8, 500 mM NaCl with protease and phosphatase inhibitors and 100 U/mL RNaseIN Plus); and 3) 2 minutes with rotation in Wash buffer 3 (SG IP buffer with 2 M urea). After removing the wash buffer, dry beads were stored at −80°C until use for mass spectrometry. For immunoblotting, beads were resuspended in 1x Laemmli buffer containing 100 mM DTT and boiled at 95°C for 10 minutes.

### Tandem-mass tag (TMT) isobaric labeling and mass spectrometry-based proteomics

#### Protein digestion and TMT labeling

Seventy-five µg of RIPA lysate (prepared as outlined earlier) or Dynabeads with bound SG cores were submitted to the University of Michigan Proteomics Resource Facility. Samples were proteolyzed and labeled with TMTpro 16-plex reagents (Thermo Fisher) following manufacturer’s instructions. Briefly, for RIPA lysate, upon reduction (5 mM DTT for 30 min at 45°C) and alkylation of cysteines (15 mM 2-chloroacetamide for 30 min at RT), the proteins were precipitated by adding 6 volumes of ice-cold acetone followed by overnight incubation at −20°C. The precipitate was spun down, and the pellet was allowed to air dry. The pellet was resuspended in 0.1M TEAB and overnight digestion with trypsin/Lys-C mix (Promega) at a 1:25 ratio of trypsin to protein at 37° C was performed with constant mixing using a thermomixer. For SG cores on Dynabeads, upon reduction and alkylation as described above, on-bead digestion with trypsin/Lys-C mix (∼1 mg) was carried out. Peptides desalted using SepPak C18 cartridge following manufacturer’s protocol (Waters; WAT023501), dried completely using vacufuge and reconstituted in 0.1M TEAB in preparation for TMT labeling. The TMTpro 16-plex reagents were dissolved in 20 µl of anhydrous acetonitrile and labeling was performed by transferring the entire digest to TMTpro reagent vial and incubating at room temperature for 1 h. Reaction was quenched by adding 8 µl of 5% hydroxyl amine and further 15 min incubation. Labeled samples were mixed and dried using a vacufuge. Combined lysate sample (∼300 mg) was fractionated into 12 concatenated fractions using high-pH reversed phase chromatography (Zorbax 300Extend-C18, 2.1mm x 150 mm column on an Agilent 1260 Infinity II HPLC system over a continuous gradient of 3-90% 20 mM ammonium bicarbonate/acetonitrile buffer system in 90 min;). For on-bead digested peptides, an offline fractionation of the combined sample into 8 fractions was performed using high pH reversed-phase peptide fractionation kit (Thermo Fisher) according to the manufacturer’s protocol. Fractions were dried and reconstituted in 9 µl of 0.1% formic acid/2% acetonitrile in preparation for LC-MS/MS analysis. The samples were labeled with TMT channels as follows:

| Channel | Sample for whole cell lysate proteomics | Sample for SG IP (on Dynabeads) proteomics |
| --- | --- | --- |
| 126 | WT HEK293 replicate 1 | NTC HeLa replicate 1 |
| 127N | WT HEK293 replicate 2 | RTL8 KO HeLa replicate 1 |
| 128N | WT HEK293 replicate 3 | PEG10 KO HeLa replicate 1 |
| 129N | NTC clone 1 HEK293 replicate 1 | UBQLN2 KO HeLa replicate 1 |
| 130N | NTC clone 1 HEK293 replicate 2 | NTC HeLa replicate 2 |
| 131N | NTC clone 1 HEK293 replicate 3 | RTL8 KO HeLa replicate 2 |
| 132N | NTC clone 2 HEK293 replicate 1 | PEG10 KO HeLa replicate 2 |
| 133N | NTC clone 2 HEK293 replicate 2 | UBQLN2 KO HeLa replicate 2 |
| 134N | NTC clone 2 HEK293 replicate 3 | NTC HeLa replicate 3 |
| 127C | RTL8 KO clone 1 HEK293 replicate 1 | RTL8 KO HeLa replicate 3 |
| 128C | RTL8 KO clone 1 HEK293 replicate 2 | PEG10 KO HeLa replicate 3 |
| 129C | RTL8 KO clone 1 HEK293 replicate 3 | UBQLN2 KO HeLa replicate 3 |
| 130C | RTL8 KO clone 2 HEK293 replicate 1 | NTC HeLa replicate 4 |
| 131C | RTL8 KO clone 2 HEK293 replicate 2 | RTL8 KO HeLa replicate 4 |
| 132C | RTL8 KO clone 2 HEK293 replicate 3 | PEG10 KO HeLa replicate 4 |
| 133C | N/A | UBQLN2 KO HeLa replicate 4 |

#### Liquid chromatography-mass spectrometry analysis (LC-multinotch MS3)

To obtain superior quantitation accuracy, multinotch-MS3^79^ was employed which minimizes the reporter ion ratio distortion resulting from fragmentation of co-isolated peptides during MS analysis. Orbitrap Fusion (Thermo Fisher) with RSLC Ultimate 3000 nano-UPLC (Dionex) or Orbitrap Ascend Tribrid equipped with FAIMS source (Thermo Fisher) with Vanquish Neo UHPLC was used to acquire the whole cell lysate or SG IP data, respectively. 2 µL of the whole cell lysate sample was resolved on a PepMap RSLC C18 column (75 µm i.d.x 50 cm; Thermo Scientific) at the flowrate of 300 nl/min using 0.1% formic acid/acetonitrile gradient system (2-22% acetonitrile in 150 min;22-32% acetonitrile in 40 min; 20 min wash at 90% followed by 50 min re-equilibration). 2 µL of the SG IP sample was resolved on an Easy-Spray PepMap Neo column (75 µm i.d. x 50 cm; Thermo Scientific) at the flowrate of 300 nl/min using 0.1% formic acid/acetonitrile gradient system (3-19% acetonitrile in 72 min;19-29% acetonitrile in 28 min; 29-41% in 20 min followed by 10 min column wash at 95% acetonitrile and re-equilibration). Samples were then directly sprayed onto the mass spectrometer using EasySpray source (Thermo Fisher). FAIMS source was operated in standard resolution mode, with a nitrogen gas flow of 4.2 L/min, and inner and outer electrode temperature of 100 °C and dispersion voltage or −5000 V. Two compensation voltages (CVs) of −45 and −65 V, 1.5 seconds per CV, were employed to select ions that enter the mass spectrometer for MS1 scan and MS/MS cycles. For whole cell lysate samples, mass spectrometer was set to collect one MS1 scan (Orbitrap; 120K resolution; AGC target 2×10^5^; max IT 100 ms) followed by data-dependent, “Top Speed” (3 seconds) MS2 scans (collision induced dissociation; ion trap; NCE 35; AGC 5×10^3^; max IT 100 ms). For SG IP samples, mass spectrometer was set to collect MS1 scan (Orbitrap; 400-1600 m/z; 120K resolution; AGC target of 100%; max IT in Auto) following which precursor ions with charge states of 2-6 were isolated by quadrupole mass filter at 0.7 m/z width and fragmented by collision induced dissociation in ion trap (NCE 30%; normalized AGC target of 100%; max IT 35 ms). For multinotch-MS3, top 10 precursors from each MS2 were fragmented by HCD followed by Orbitrap analysis with settings as follows: NCE 55; 60K resolution; AGC 5×10^4^; max IT 120 ms, 100-500 m/z scan range or NCE 55; 45K resolution; normalized AGC target of 200%; max IT 200 ms, 100-500 m/z scan range or for whole cell lysates or SG IP samples, respectively.

#### Data Analysis

Proteome Discoverer (Thermo Fisher) was used for data analysis. MS2 spectra were searched against SwissProt human protein database (Whole cell lysate: 20291 entries, downloaded on 12/31/21; SG IP: 20352 entries, downloaded on 11/2/23) using the following search parameters: MS1 and MS2 tolerance were set to 10 ppm and 0.6 Da, respectively; carbamidomethylation of cysteines (57.02146 Da) and TMT labeling of lysine and N-termini of peptides (304.207 Da) were considered static modifications; oxidation of methionine (15.9949 Da) and deamidation of asparagine and glutamine (0.98401 Da) were considered variable. Identified proteins and peptides were filtered to retain only those that passed ≤1% FDR threshold. Quantitation was performed using high-quality MS3 spectra (Signal-to-noise ratio of 12 (for whole cell lysates) or 15 (for SG IP) and 60% isolation interference). For whole cell lysate proteomics, peptide usage was set to Unique to capture peptides unique to PEG10 gag-pol. A new control group was created by combining samples from WT and NTC HEK293 cells. For SG IP proteomics, PSM threshold was set to 5 to exclude low abundance hits. T-tests were performed with null hypothesis that log_2_ fold change versus NTC or control group was 1. Volcano plots and heatmaps were generated on RStudio (Posit). Volcano plots were obtained by plotting log_10_ (adjusted p-value) to log_2_ fold change versus NTC. SG proteins were identified using the RNA granule database.^80^ For heatmaps, a list of PEG10 interactors was made using IP/MS data from Pandya et al., 2021 and the BioGrid database.^21,81^ EV proteins were obtained from Xu et al., 2016.^82^ Raw mass spec data and R code have been deposited.

### Biotinylated isoxazole (B-isox) precipitation

B-isox precipitation was performed as described previously.^83,84^ Cells were lysed in EE buffer (50 mM HEPES pH 7.5, 150 mM NaCl, 0.1% NP40, 10% glycerol, 1 mM DTT, 1 mM EDTA, 2.5 mM EGTA and 1U/μL RNaseIN Plus supplemented with protease and phosphatase inhibitors) with rotation for 20 minutes at 4°C and centrifuged at 16,000xg for 15 minutes at 4°C. A small fraction of the supernatant was saved for input. The remaining supernatant was divided in half: B-isox dissolved in DMSO was added to one half at a final concentration of 0.1 mM and an equivalent volume of DMSO was added to the other. Lysates were incubated with rotation for 1.5 h at 4°C and centrifuged at 10,000xg for 10 minutes at 4°C. The supernatants were harvested to which Laemmli buffer was added and boiled for 10 minutes. The pellets were washed twice by resuspending in ice-cold EE buffer, vortexing, incubating on ice for 10 minutes followed by centrifuging at 10,000xg for 10 minutes at 4°C. They were resuspended in 1x Laemmli buffer with 100 mM DTT and boiled for 10 minutes.

### VLP isolation and protease protection assay

VLPs were isolated as described previously^33^. HeLa PEG10 KO cells were grown on 15 cm dishes and transfected with pCMV-PEG10 gag-pol-Dendra2 using Lipofectamine 3000. 72 h after transfection, the media was collected and precleared by spinning at 2700 xg for 15 minutes at 4°C, then 10,000 xg for 10 minutes at 4°C. The precleared media was then layered dropwise onto a 30% (w/v) sucrose cushion in an ultracentrifuge tube (Beckman Coulter). Ultracentrifugation was done at 134,000xg for 4h at 4°C using an SW 41 Ti rotor (Beckman Coulter) in an Optima XE-90 ultracentrifuge (Beckman Coulter). The resulting VLP pellet was resuspended either in PBS for protease protection assays or PBS with 10% sucrose for cryo-CLEM, fluorescence SEC and negative stain TEM.

Protease protection assay was performed as outlined previously.^32^ VLP suspensions were split into three tubes. Tube 1: 20 µL VLP suspension + 7 µL water, tube 2: 20 µL VLP suspension + 6 µL water + 1 µL Proteinase K (200 µg/mL, NEB), and tube 3: 20 µL VLP suspension + 3 µL water + 3 µL 10% Triton X-100 + 1 µL Proteinase K (200 µg/mL). After addition of Proteinase K, samples were incubated for 10 minutes at RT, following which 1 µL of 30 mM PMSF (in DMSO) was added and incubated for 10 minutes at RT. Laemmli buffer was added to each tube to a final concentration of 1x and boiled for 10 minutes.

### Negative stain TEM

Negative staining was carried out with 0.75% (w/v) uranyl formate (Electron Microscopy Sciences) as described previously.^85^ Briefly, 300-mesh copper grids with carbon-coated film were glow discharged for 60 s at 10 mA using an EasiGlow system (Pelco). 3.5 μL of crude VLP suspension was pipetted onto the grids and incubated for 1 minute at room temperature. Two wash cycles were performed with 30 μL of deionized water, followed by one wash with 30 μL of uranyl formate, and finally stained with 30 μL of uranyl formate for 1 minute. Prepared grids were imaged on a Morgagni transmission electron microscope (FEI) equipped with a field emission gun operating at an acceleration voltage of 100 keV set to a nominal magnification of 22,000× (2.5 Å/pixel).

### Fluorescence size exclusion chromatography (SEC)

Crude VLP suspensions were filtered through a 0.45 μm filter (Corning) and analyzed on a Superose 6 Increase 10/300 GL column (Cytiva) pre-equilibrated with PBS. Dendra2 fluorescence in the eluate was monitored using a fluorescence detector (Shimadzu Scientific Instruments) with the following specifications: Excitation - 485 nm and emission - 510 nm. Molecular weight standard peaks were determined by running a Gel Filtration Standard (BioRad), comprising peaks at 670 kDa (thyroglobulin), 158 kDa (γ-globulin), 44 kDa (ovalbumin), and 17 kDa (myoglobin), through the column separately.

### Custom cryo-confocal light electron microscopy (cryo-CLEM) workflow

#### Sample preparation for cryo-electron tomography

*Plunge-freezing VLPs:* 3 µL of crude enriched VLP resuspension was applied to Holey copper R2/2 grids (Quantifoil GmBH) glow-discharged at 5mA 30s using an EasiGlow system (Pelco). The grids were plunge-frozen using a Vitrobot Mark IV (Thermo Fisher) with the following specifications: *blot time* - 2.5s, *blot force* - 1, *temperature* - 4°C, and *humidity* - 90%. The plunge-frozen grids were clipped into AutoGrids (Thermo Fisher) and stored in liquid nitrogen until further use.

#### Plunge-freezing HeLa cells

HeLa cells were vitrified as described previously.^86^ Briefly, gold mesh grids with holey R1/4 SiO_2_ film (Quantifoil Micro Tools GmbH) were glow discharged for 60s at 15 mA using an EasiGlow system. Subsequently, the grids were treated with PEG-SVA (Laysan Bio Inc.) and UV-sensitive photoinitiator (PLPP, Alvéole Lab) solutions before ablating patterns onto the film with a PRIMO digital-micromirror device (Alvéole Lab). Micropatterned grids were transferred to a 35 mm glass bottom dish (MatTek) onto which PEG10 KO G3BP1-mScarlet-I HeLa cells were seeded. Cells were transfected 48 h later with 1.5 μg of pCMV-PEG10 gag-pol-Dendra2 using Lipofectamine 3000. They were stressed with sodium arsenite 24 h post transection, following which 4 μL of 1:125 diluted crimson FluoSpheres (Thermo Fisher) were added before back-side blotting and plunge freezing into liquid ethane at −185 °C using a GP2 (Leica Microsystems). The plunge-frozen grids were clipped into FIB-AutoGrids (Thermo Fisher) and were stored in liquid nitrogen until further use.

#### Cryo-fluorescence microscopy

Positioning and fluorescence of VLPs/cells on grids was assessed in widefield mode on a Stellaris-5 cryo-confocal laser scanning microscope equipped with a cryo-stage and x50/0.9 numerical aperture objective (Leica Microsystems). Z-stacks of desired regions were acquired in confocal-scan mode with a 490 nm laser (505-540 nm emission), 570 nm laser (585-610 nm emission), 625 nm laser (635-650 nm emission), 0.5 µm spacing spanning ± 7 µm of the focus plane and an x/y pixel size of 110 nm. Additionally, a reflection image was acquired at each Z plane to position the PEG10-Dendra2 or FluoSpheres signal relative to the holey-carbon film for targeted focused ion beam milling via 3D Correlation Toolbox (3DCT) and/or tilt-series data acquisition. MIPs of Z-stacks were generated on LAS X.

#### Cryo-focused ion beam (cryo-FIB) milling

Screened grids were transferred into a dual-beam Aquilos 2 FIB-SEM (Thermo Fisher) maintained at cryogenic temperature. Grids were coated with a layer of organometallic platinum using a gas injection system for 30 s and then sputter-coated with inorganic platinum to form a protective/conductive surface for milling. Subsequently, correlation between cryo-confocal, SEM, FIB, and TEM images was performed on the cross-platform correlative software Maps 3 (Thermo Fisher) and 3DCT. First, a three-point correlation between widefield and SEM (100x magnification at 50 kV) images was used for initial XY grid navigation in search of ROIs captured in cryo-confocal. Next, SEM (1,500x magnification at 50 kV) and FIB (2,500x magnification at 10 pA) images were captured at a milling angle of 15° to screen for ice contamination and distinguishable FluoSpheres on the grid’s carbon film. In the 3DCT user interface, corresponding FluoSpheres were identified in the confocal fluorescence Z-stack and the ion beam image. They were used to determine the transformation parameters for other channels. Transformed MIP for PEG10-Dendra2 and G3BP1-mScarlet-I guided initial cuts with the FIB to make tension-relief trenches. Subsequently, 5µm thick lamella was milled with a beam current of 1 nA at a milling angle of 15°, and 1µm lamella with 100-500 pA beam currents. Milling progress and lamella integrity were monitored via SEM imaging. Upon completion of all rough lamella on a grid, each lamella was revisited and polished to a thickness of ∼200 nm with the FIB at a beam current of 50 pA.

#### Tilt-series data acquisition, processing, and segmentation

Plunge-frozen grids containing VLPs or milled cells were imaged on a 300 kV Titan Krios G4i transmission electron microscope (Thermo Fisher), equipped with a K3 direct electron detector and an imaging filter (Gatan Inc.) operated in counting mode. Dose-symmetric tilt-series (as outlined in ^87^) were collected from –60° to +60°, starting at 0° at a 3° interval, at correlated areas using SerialEM software.^73^ The magnification was set to a pixel size of 3.315 Å/pixel and the total dose per tilt-series was 120e^-^/Å^2^. Data were collected with a 20eV slit width. Tilt-series were aligned by patch tracking and reconstructed by weighted back projection implemented in IMOD.^70^ 15 iterations of a ‘SIRT-like’ filter were applied. Organelle and VLP membranes were segmented using MemBrain v2.^72^ Ribosome coordinates were detected by deconvolving tomograms in Warp,^74^ template matching, and using warp2dynamo to extract the coordinates. These coordinates were then used to place the ribosome density (EMD-15636). Microtubules were segmented using the cylinder correlation feature in Amira (Thermo Fisher). Intermediate filaments were automatically detected by the convolutional neural network-based segmentation implemented within Amira.

### Phylogenetic tree construction, sequence and structure alignment

A phylogenetic tree of the RTL gene family was constructed using the neighbor-joining method on MEGA^71^ with the following UniProt-annotated protein sequences: RTL1 (A6NKG5), PEG10/RTL2 (Q86TG7 isoform 1), RTL3 (Q8N8U3), RTL4 (Q6ZR62), RTL5 (Q5HYW3), RTL6 (Q6ICC9), LDOC1/RTL7 (O95751), RTL8A (Q9BWD3), RTL8B (Q17RB0), RTL8C (A6ZKI3), RTL9 (Q8NET4), and Bop/RTL10 (Q7L3V2). Sequences were aligned using the MUSCLE algorithm. Then, ambiguous positions for each sequence pair were removed using the pairwise deletion method prior to employing the Poisson correction method to compute evolutionary distance.

LDOC1, RTL8A, RTL8B, and RTL8C protein sequences were aligned and visualized using Clustal Omega and ESPript 3.0,^69^ respectively. Sequence alignment, RMSD and TM scores were obtained using AlphaFold-predicted models of LDOC1 (AFO95751F1) and RTL8C (AFA6ZKI3F1) on the RCSB pairwise sequence alignment tool using the FATCAT (rigid) algorithm. The final image was generated on PyMol (Schrödinger).

### Statistical analysis

Statistical analyses were performed using GraphPad (Prism). Data from multiple replicates were used in unpaired analyses. Statistical tests performed are denoted in respective figure legends. For proteomic analyses, t-tests were performed within Proteome Discoverer (Thermo Fisher). Unless otherwise mentioned, a p-value < 0.05 was considered significant.

## RESULTS

### PEG10 is regulated by the combined actions of UBQLN2 and RTL8

The studies described here seek to understand why the related retrovirus-like proteins, RTL8A, B, and C (collectively referred to as RTL8 due to their nearly identical sequences) and PEG10 (Figure 1A and C), interact with and influence the properties of UBQLN2 in cellular proteostasis. As a first step, building on our recent discovery that RTL8 proteins are major UBQLN2 interactors,^51^ we compared the proteome of HEK293 cells knocked out for all three RTL8 proteins (RTL8 KO) to that of wild-type (WT) and non-targeting control (NTC) cells by tandem mass tag-based mass spectrometry (TMT-MS). Among the most upregulated proteins, we identified PEG10 (Figure 1B and Data S1). PEG10 undergoes ribosomal frameshifting to generate two distinct isoforms, the shorter gag and longer gag-pol isoforms (Figure 1D).^88^ Loss of RTL8 selectively increases the gag-pol isoform (Figure 1E, S1A and S1D). This increase is not due to a change in UBQLN2 levels (Figure S1A). Moreover, levels of two canonical UBQLN2 clients, TRIM32 and ATAD1,^50^ are unaffected by loss of RTL8 (Figure S1B). RTL8 KO cells also showed no change in the levels of two proteins known to modulate PEG10, the ubiquitin depeptidase USP9X and the E3 ubiquitin ligase UBE3A^21,29^ (Figure S1C). Together, these results suggest RTL8 directly regulates the cellular levels of PEG10 gag-pol.

To confirm this finding, we performed siRNA knockdown of RTL8 or UBQLN2 in a panel of human cell lines (Figures 1F and S1E-G). UBQLN2 knockdown increased PEG10 gag-pol levels in all cell lines, and RTL8 knockdown caused an even greater increase in gag-pol without altering UBQLN2 levels. For simplicity, we used RTL8C as a representative for all RTL8 proteins in subsequent experiments. Transiently adding back RTL8C to RTL8 KO cells lowered PEG10 gag-pol without altering UBQLN2 (Figure 1G), and prolonged RTL8C expression reduced PEG10 gag-pol even further (Figure S1H). This action of RTL8 does not extend to its closest paralog, leucine zipper downregulated in cancer 1 (LDOC1, a.k.a. RTL7) (Figure 1C, D and G), which is also a PEG10 interactor.^16,19^ Since the only major differences between LDOC1 and RTL8C are in their N-termini (Figure S2A), we tested whether the unique N-terminus of RTL8 was essential for PEG10 regulation. Indeed, RTL8C lacking N-terminal residues Asp2 to Asn25 (referred to as RTL8C ΔN) was unable to reduce PEG10 gag-pol levels. Thus, PEG10 gag-pol is regulated by RTL8 in a manner dependent on its unique N-terminus.

We next sought to determine whether the ability of RTL8 to regulate PEG10 gag-pol requires UBQLN2. To do so, we examined HEK293 cells knocked out for all three brain-expressed UBQLNs, UBQLNs 1, 2 and 4 (UBQLN TKO cells).^67^ Because RTL8 is normally stabilized by UBQLN2,^50,51^ UBQLN TKO cells also have markedly reduced RTL8 levels, which is restored by reintroducing UBQLN2 (Figure S2B). Transiently expressing UBQLN2 alone reduced PEG10 gag-pol and co-expressing UBQLN2 with RTL8C did not further reduce PEG10 gag-pol. These results indicate that RTL8 is unable to regulate PEG10 gag-pol without UBQLN2. To test whether RTL8 is essential for UBQLN2-mediated clearance of PEG10, we transiently expressed UBQLN2 in UBQLN TKO cells that were simultaneously treated with siRNA targeting RTL8. After RTL8 knockdown, UBQLN2 no longer reduces PEG10 gag-pol levels but when siRNA-resistant RTL8C is added back, UBQLN2 once again decreases PEG10 gag-pol (Figure S2C). Thus, RTL8 is an essential partner for UBQLN2-mediated regulation of PEG10.

### PEG10-UBQLN2 interactions require the N-terminus of RTL8

Because UBQLN2 requires RTL8 to regulate PEG10, we hypothesized that RTL8 enables UBQLN2 interactions with PEG10. To test this, we performed co-immunoprecipitation (co-IP) assays in which endogenous UBQLN2 was pulled down from NTC or RTL8 KO cells in the presence or absence of the reversible crosslinker, DTBP (Figure 2A). RTL8 and PEG10 gag-pol co-precipitated with UBQLN2, and the amount of co-precipitating PEG10 gag-pol was much lower in RTL8 KO cells (to a lesser extent, the shorter PEG10 gag isoform also co-precipitated with UBQLN2 in an RTL8-dependent manner). The use of crosslinker to stabilize interactions did not affect PEG10 co-IP with UBQLN2 but did, in contrast, lead to a significant increase in RTL8 co-IP, suggesting that the UBQLN2-RTL8 interaction is of lower affinity.

**Figure 2:**
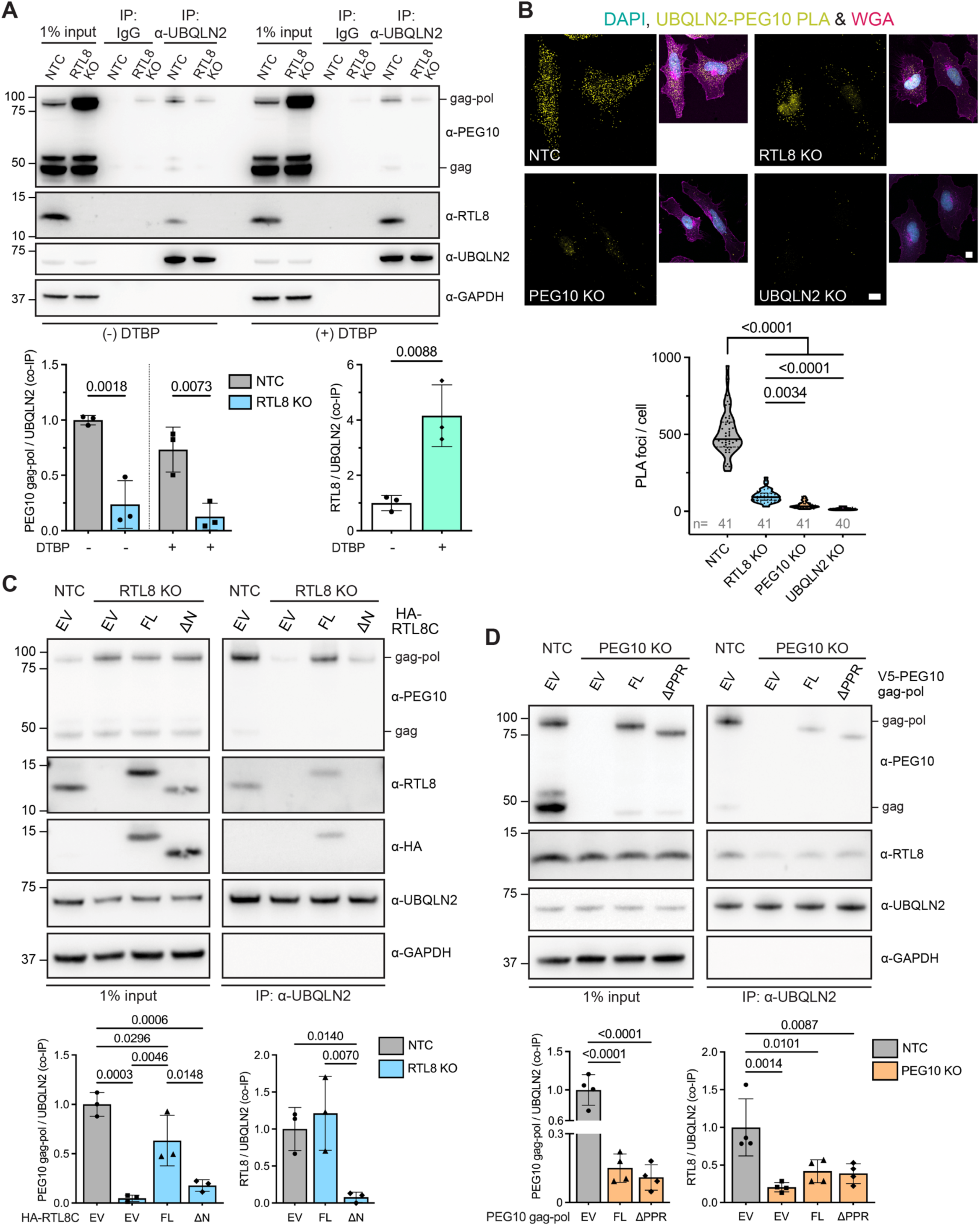
PEG10-UBQLN2 interactions require the N-terminus of RTL8. **A)** Lysates from NTC or RTL8 KO HEK293 cells were IPed with control or α-UBQLN2 antibody in the absence or presence of the crosslinker DTBP. The amount of PEG10 or RTL8 co-IPed, relative to IPed UBQLN2 and, normalized to NTC HEK293 without DTBP addition, are shown below. **B)** PLA assessing interaction between endogenous UBQLN2 and PEG10 in the indicated HeLa cell lines. Nuclei and cell membranes are labeled with DAPI and wheat germ agglutinin (WGA), respectively. Quantification of PLA foci/cell is shown below. Number of individual cells quantified per line from 2 independent replicates is indicated. Scale bars: 10 μm. Lysates from **C)** NTC or RTL8 KO HeLa cells expressing full-length (FL) HA-RTL8C, HA-RTL8C ΔN or empty vector, or **D)** NTC or PEG10 KO HeLa cells expressing FL V5-PEG10 gag-pol, V5-PEG10 gag-pol ΔPPR or empty vector, were IPed with α-UBQLN2 Ig. Amount of co-IPed PEG10 or RTL8, relative to IPed UBQLN2 and normalized to NTC HeLa is shown below. Also, see Figure S3. In (A-D), data were analyzed using one-way ANOVA with Tukey’s multiple comparison test. In (A, C, and D), data represent means ± SD (N=3), GAPDH = loading control.

Next, we performed a proximity ligation assay (PLA) to assess interactions between endogenous UBQLN2 and PEG10 in HeLa cells; control PLA experiments were performed in cells lacking RTL8, PEG10, or UBQLN2. NTC HeLa cells contained far more PLA foci than RTL8 KO cells, confirming that RTL8 mediates interactions between UBQLN2 and PEG10 (Figure 2B). Because RTL8 KO cells had more PLA foci than either PEG10 KO or UBQLN2 KO cells, we conclude that UBQLN2 and PEG10 still interact in the absence of RTL8, but less robustly.

Because the N-terminus of RTL8 is required to reduce PEG10 gag-pol levels, we surmised that this domain likely facilitates UBQLN2-PEG10 gag-pol interactions. To test this, we pulled down UBQLN2 from RTL8 KO cells expressing full-length RTL8C or RTL8C ΔN (Figure 2C). Full-length RTL8C co-precipitated with UBQLN2 and partially restored UBQLN2-PEG10 gag-pol interactions, whereas RTL8C ΔN did not. Likewise, only full-length RTL8C co-precipitated with PEG10 and rescued UBQLN2 interactions with PEG10 gag-pol (Figure S3A). Thus, the N-terminus of RTL8 enables separate interactions with UBQLN2 and PEG10 and thereby facilitates interactions between UBQLN2 and PEG10.

PEG10 gag-pol contains an intrinsically disordered C-terminal polyproline region (PPR) that is important for regulation by UBQLN2.^22^ Accordingly, we next tested whether the PPR is required for PEG10 interactions with RTL8 or UBQLN2. In PEG10 KO cells, transiently expressed full-length or ΔPPR PEG10 gag-pol both co-precipitated similarly with UBQLN2 (Figure 2D). We confirmed these results in reciprocal IP experiments, in which we pulled down PEG10 gag-pol and assessed coprecipitation of UBQLN2 (Figure S3B). Thus, PEG10 gag-pol’s PPR is not required to interact with either UBQLN2 or RTL8.

To understand the order of interactions between the three proteins, we pulled down UBQLN2 from PEG10 KO cells or PEG10 from UBQLN2 KO cells, then compared the amount of co-IPed RTL8, normalized to input (Figure S3C). In the absence of PEG10, much less RTL8 co-precipitated with UBQLN2. In the absence of UBQLN2, however, RTL8 continued to co-precipitate with PEG10 gag-pol. We also noted that the loss of PEG10 markedly reduced RTL8 interactions with UBQLN2 (Figure 2D). Taken together, these data suggest that RTL8 and PEG10 gag-pol likely interact first and then recruit UBQLN2.

### PEG10-UBQLN2 interactions are preserved in SGs

To understand the biological implications of the interactions between PEG10, RTL8, and UBQLN2, we chose to study them in the context of SGs (Figure 3A). SGs are ribonucleoprotein (RNP) condensates that form as a result of stress-induced stalling of translation.^52,54^ While RTL8 is not known to be a SG component, both PEG10 and UBQLN2 have been shown to localize to SGs.^21,38,89^ Accordingly, we hypothesized that PEG10-UBQLN2 interactions would persist within SGs. To test this, we performed PLA on cells treated with either the oxidative stressor sodium arsenite or heat shock to induce SG formation, which was visualized by immunostaining for the SG core protein G3BP1 (Figure 3B). PLA foci density was significantly greater within G3BP1-positive SGs, indicating that PEG10-UBQLN2 interactions are enriched within SGs.

**Figure 3:**
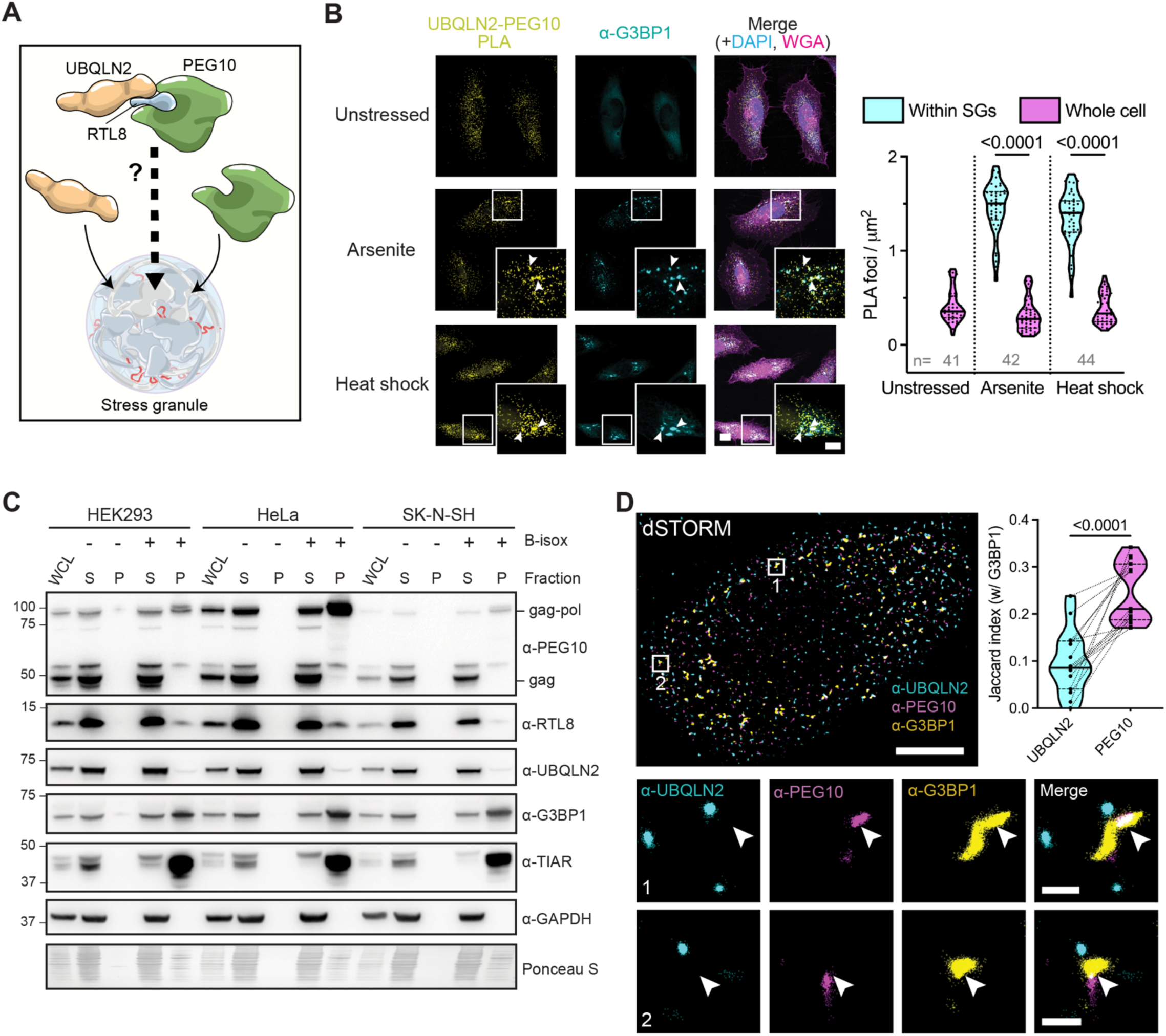
PEG10-UBQLN2 interactions are preserved in SGs. **A)** Schematic depicting possible SG localization of the UBQLN2-RTL8-PEG10 complex given the known SG localization of UBQLN2 and PEG10. **B)** PLA assessing the interaction between endogenous UBQLN2 and PEG10 in HeLa cells induced to form SGs by sodium arsenite or heat shock. Nuclei and cell membranes are labeled with DAPI and WGA, respectively, and arrowheads in insets mark SGs stained with G3BP1. Quantification of PLA foci/μm^2^ within SGs and in the whole cell is shown below. Number of individual cells quantified per condition from 2 independent replicates is indicated. One-way ANOVA with Tukey’s multiple comparison test was performed. Scale bars: 10 µm and 5 µm for insets. Also, see Figure S4B. **C)** Biotinylated isoxazole (B-isox)-mediated precipitation of RNA granule proteins from three human cell lines. WCL = whole cell lysate before B-isox addition, S = supernatant, and P = pellet. G3BP1 and TIAR are positive controls, GAPDH is a negative control. Ponceau S was used to normalize loading. **D)** 3-color dSTORM of HeLa cells treated with sodium arsenite. Numbers identify the zoomed regions shown below. Arrowheads mark regions of overlap between PEG10 and G3BP1. Jaccard index between G3BP1 clusters and UBQLN2 or PEG10 is shown at top right. Dotted lines connect data points from an individual cell. An unpaired t-test was performed. N=15 cells across 3 independent replicates. Scale bars: 10 µm and 0.5 µm for zoomed regions.

Proteins that have a tendency to form biomolecular condensates like SGs can be captured in biotinylated isoxazole (B-isox)—mediated precipitation assays.^83,84,90,91^ Since UBQLN2 and PEG10 interactions are enriched in SGs and are mediated by RTL8, we reasoned that all three proteins should be enriched in B-isox precipitates. Indeed, across multiple tested human cell lines, we observed co-precipitation of endogenous RTL8, UBQLN2 and PEG10 gag-pol (but not gag), upon addition of B-isox (Figure 3C). Of the three proteins, PEG10 gag-pol was most enriched in the B-isox pellet, mirroring the behavior of SG core proteins, G3BP1 and TIAR.^92^

We then spatially mapped the distribution of UBQLN2 and PEG10 in SGs using direct stochastic optical reconstruction microscopy (dSTORM), a technique used to visualize SG ultrastructure.^92,93^ As measured by the Jaccard index, PEG10 clusters overlapped more with G3BP1 clusters than did UBQLN2 clusters, which also were near to and sometimes overlapped with G3BP1 (Figure 3D). This result suggests that PEG10 and UBQLN2 tend to localize to distinct subregions in SGs and that PEG10, unlike UBQLN2, is directly embedded within SG cores. These findings are consistent with a recent study identifying PEG10 as a core component of stress-induced condensates.^94^

### PEG10 recruits UBQLN2 to SGs in an RTL8-dependent manner

Because PEG10-UBQLN2 interactions require RTL8, we sought to understand whether the loss of RTL8 affects recruitment of UBQLN2 or PEG10 to SGs. In both HeLa and HEK293 cells, UBQLN2 and PEG10 readily localized to SGs after treatment with sodium arsenite (Figure 4A and S4A). In RTL8 KO cells, however, UBQLN2 failed to localize to SGs, whereas PEG10 still did. Similarly, in PEG10 KO cells UBQLN2 no longer localized to SGs, illustrating that UBQLN2 requires both RTL8 and PEG10 for recruitment to SGs. In contrast, PEG10 does not require UBQLN2 or RTL8 for SG localization and actually becomes enriched in SGs when either protein is absent.

**Figure 4:**
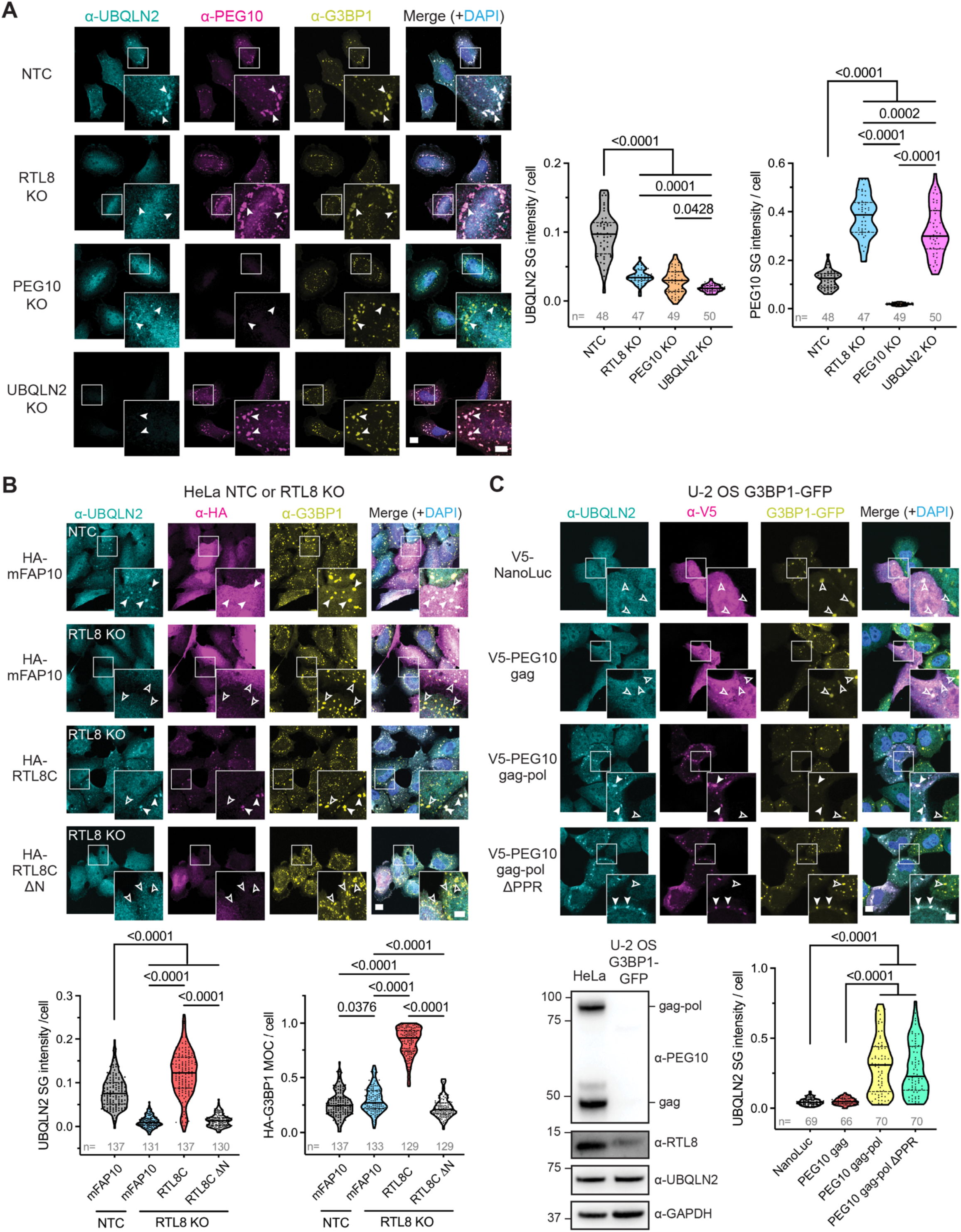
PEG10 gag-pol recruits UBQLN2 to SGs in an RTL8-dependent manner. **A)** After treatment with sodium arsenite, the indicated HeLa cell lines were immunostained for endogenous UBQLN2, PEG10, and the SG marker G3BP1. Arrowheads in insets mark SGs. UBQLN2 or PEG10 intensity within SGs was quantified and is shown to the right. Also, see Figures S4 and S5A. **B)** NTC and RTL8 KO HeLa cells expressing HA-RTL8C, HA-RTL8C ΔN, or control protein (HA-mFAP10), treated with sodium arsenite. UBQLN2 intensity within SGs is shown at bottom left. The Mander’s overlap coefficient (MOC) between HA and G3BP1 is shown at bottom right. **C)** U-2 OS G3BP1-GFP cells expressing V5-PEG10 gag, gag-pol, gag-pol ΔPPR, or control protein (V5-Nanoluciferase) treated with sodium arsenite. Immunoblot of lysates from HeLa and U-2 OS G3BP1-GFP cells is shown at bottom left. UBQLN2 intensity within SGs is shown at bottom right. Also, see Figure S5B. In (A-C), nuclei are labeled with DAPI. Scale bars: 10 µm and 5 µm for insets. In (B, C), filled arrowheads in insets indicates SGs with UBQLN2, unfilled arrows indicate SG without UBQLN2. In all panels, one-way ANOVA with Tukey’s multiple comparison test was performed. The number of individual cells quantified per line/plasmid from 3 independent replicates is indicated.

Similar results were obtained across a panel of stressors including heat shock, hyperosmotic stress and dsRNA stress, indicating that UBQLN2’s failure to localize to SGs in RTL8 and PEG10 KO cells is not stress-specific (Figures S4B-E). Likewise, the increased localization of PEG10 to SGs in the absence of RTL8 or UBQLN2 occurred with all stressors. To rule out clonal artifacts, we stressed WT HeLa cells in which RTL8, PEG10 or UBQLN2 was knocked down with siRNA (Figure S5A). Again, UBQLN2 failed to localize to SGs when RTL8 or PEG10 was transiently knocked down.

In RTL8 KO cells, expression of full-length RTL8C rescued UBQLN2 localization to SGs whereas neither RTL8C ΔN nor a similarly-sized control protein (mFAP10)^62^ did (Figure 4B). Moreover, full-length RTL8C but not RTL8C ΔN localized to SGs. These data point to the requirement of functional interactions between UBQLN2 and PEG10, mediated by RTL8, in order for UBQLN2 to localize to SGs. These data also suggest that RTL8 is a novel SG protein whose N-terminus mediates its SG recruitment.

We next determined which PEG10 isoform was responsible for shuttling UBQLN2 to SGs. An earlier study showed that only the gag-pol isoform localizes to SGs.^21^ We generated a HeLa cell line specifically knocked out for PEG10 gag-pol while maintaining levels of PEG10 gag (Figure S5B). UBQLN2 did not localize to SGs in these cells, confirming that gag-pol, but not gag, is required for UBQLN2 localization. To answer whether PEG10 gag-pol is sufficient to recruit UBQLN2 to SGs, we expressed each isoform separately in U-2 OS G3BP1-GFP cells, which naturally lack PEG10 but express UBQLN2 and RTL8 (Figure 4C). Whereas expressing gag-pol resulted in robust UBQLN2 localization to SGs, expressing gag or a control protein (NanoLuciferase) did not. The PPR of gag-pol also proved dispensable for UBQLN2 recruitment to SGs. Taken together these results indicate that PEG10 gag-pol recruits UBQLN2 to SGs in an RTL8-dependent manner.

### Depletion of UBQLN2, RTL8 or PEG10 affects SG formation and disassembly

Changes to the dynamics of SG formation and dissolution are implicated in the pathology of diseases such as ALS.^52,59,95^ UBQLN2 was previously shown to limit SG formation, whereas the effects of PEG10 and RTL8 on SG dynamics are unknown.^89^ Depletion of RTL8 or PEG10 generates an intriguing cellular scenario: UBQLN2 is expressed at near physiological levels yet unable to localize to SGs. We hypothesized that RTL8 and PEG10 KO cells, like UBQLN2 KO cells, would have a faster rate of SG formation. To test this, we tagged endogenous G3BP1 with mScarlet-I in our series of HeLa KO lines and monitored SG formation (Figure 5A and S6A). As expected, UBQLN2 KO cells formed SGs more rapidly than NTC, RTL8 KO or PEG10 KO cells, all of which showed similar rates of SG formation (Figure 5B and Movie S1). However, the loss of any of the three proteins led to smaller SGs whose G3BP1 intensity was proportional to their size (Figure S6B). Thus, loss of RTL8 or PEG10 (and therefore loss of UBQLN2 localization to SGs) does not affect SG formation but does influence SG size.

**Figure 5:**
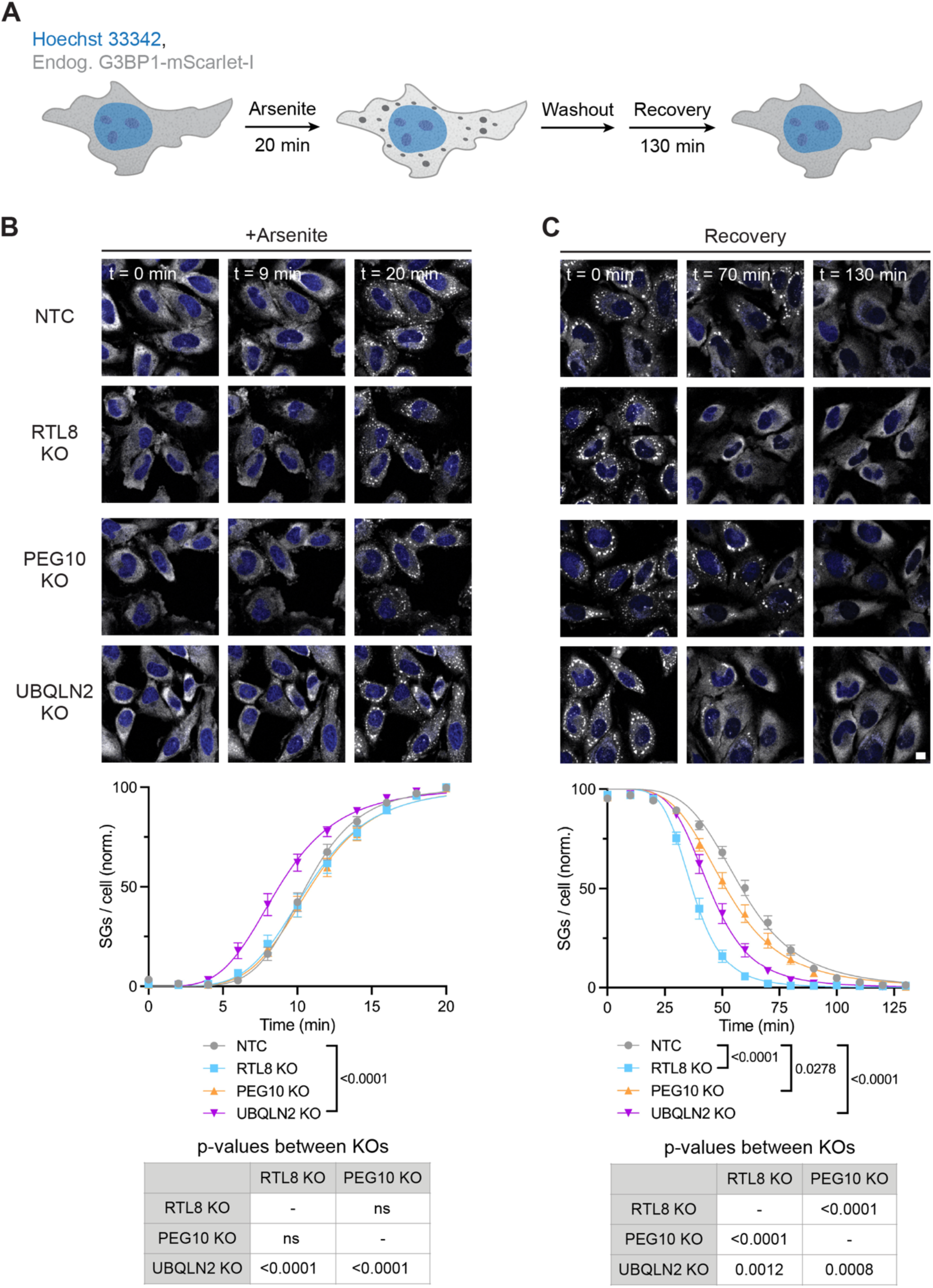
Depletion of PEG10, RTL8, or UBQLN2 alters SG dynamics. **A)** Schematic of live-cell imaging experiments in HeLa cell lines expressing endogenous G3BP1-mScarlet-I. Nuclei are labeled with Hoechst 33342. Live-cell imaging showing timepoints during arsenite stress (**B**) and recovery after wash out (**C**). Shown below are SG counts per cell over time. Data represent means ± SEM. SGs/cell are normalized to the maximum number of SGs observed under each condition, fitted to a non-linear regression model and analyzed with one-way ANOVA with Tukey’s multiple comparison test. N=24 fields across 3 independent replicates. Scale bar: 10 µm. Also, see Figure S6 and Movies S1 and S2.

Since UBQLN2 is hypothesized to extract ubiquitinated proteins to promote SG disassembly,^38^ we assessed whether the loss of UBQLN2 or the retrovirus-like proteins affected SG dissolution. In HeLa cells expressing endogenously-tagged G3BP1-mScarlet-I, we measured the rate of SG dissolution during recovery from arsenite stress (Figure 5A). Unexpectedly, UBQLN2, RTL8 and PEG10 KO cells all showed faster SG dissolution rates than NTC cells, with RTL8 KO cells having the fastest rate (Figure 5C and Movie S2). These data suggest all three proteins normally slow SG dissolution. Furthermore, changes to PEG10 levels in either direction seemed to impact SG disassembly, as PEG10 KO cells and RTL8 and UBQLN2 KO cells (which express more PEG10) all resolve SGs faster. Taken together, these results indicate distinct roles for UBQLN2 in SG formation and for all three proteins in SG disassembly.

### PEG10 shuttles select EV proteins to SGs

SG composition is highly regulated and the aberrant incorporation of proteins into SGs has been linked to several human diseases.^54,96,97^ Because PEG10 and UBQLN2 are known to interact with SG components,^21,89,98^ we next determined whether the loss of PEG10, UBQLN2, or RTL8 altered SG composition. From various HeLa KO and NTC lines, we biochemically enriched SG cores and IPed G3BP1 using previously established protocols^77,78,92^ (Figure 6A). The specificity of our pulldown was confirmed by probing for the SG components ATXN2 and TIAR, which co-precipitated with G3BP1 only after arsenite treatment (Figure S7A). We then performed TMT-MS to compare the SG composition of KO lines to that of NTC HeLa cells (Data S2). After thresholding for low abundance hits, we focused on bona fide SG proteins annotated in the RNA granule database,^80^ which classifies proteins into three tiers based on experimental evidence, with tier 1 being the most validated. Across our pulldown experiments, we detected 241 tier 1, 486 tier 2, and 425 tier 3 SG proteins, together accounting for 67% of all SG proteins and representing an approximate 3.8-fold enrichment of SG proteins compared to the whole proteome (32.7% in our dataset versus 8.6% (1720 / 20,000) in the entire proteome). The loss of UBQLN2, RTL8, or PEG10 resulted in distinct changes to SG composition. As expected, we found that PEG10 was enriched in SGs from RTL8 and UBQLN2 KO cells (Figure 6B and S7B). UBQLN2 and RTL8, however, did not meet our threshold for analysis. Some proteins were commonly enriched (e.g. KHDRBS3) or depleted (e.g. OAS3) across all three cell lines. RTL8 and PEG10 KO lines also showed changes in retroelement-derived proteins (NYNRIN and L1RE1), as well as proteins associated with viral stress response (MEX3C and OAS3).

**Figure 6:**
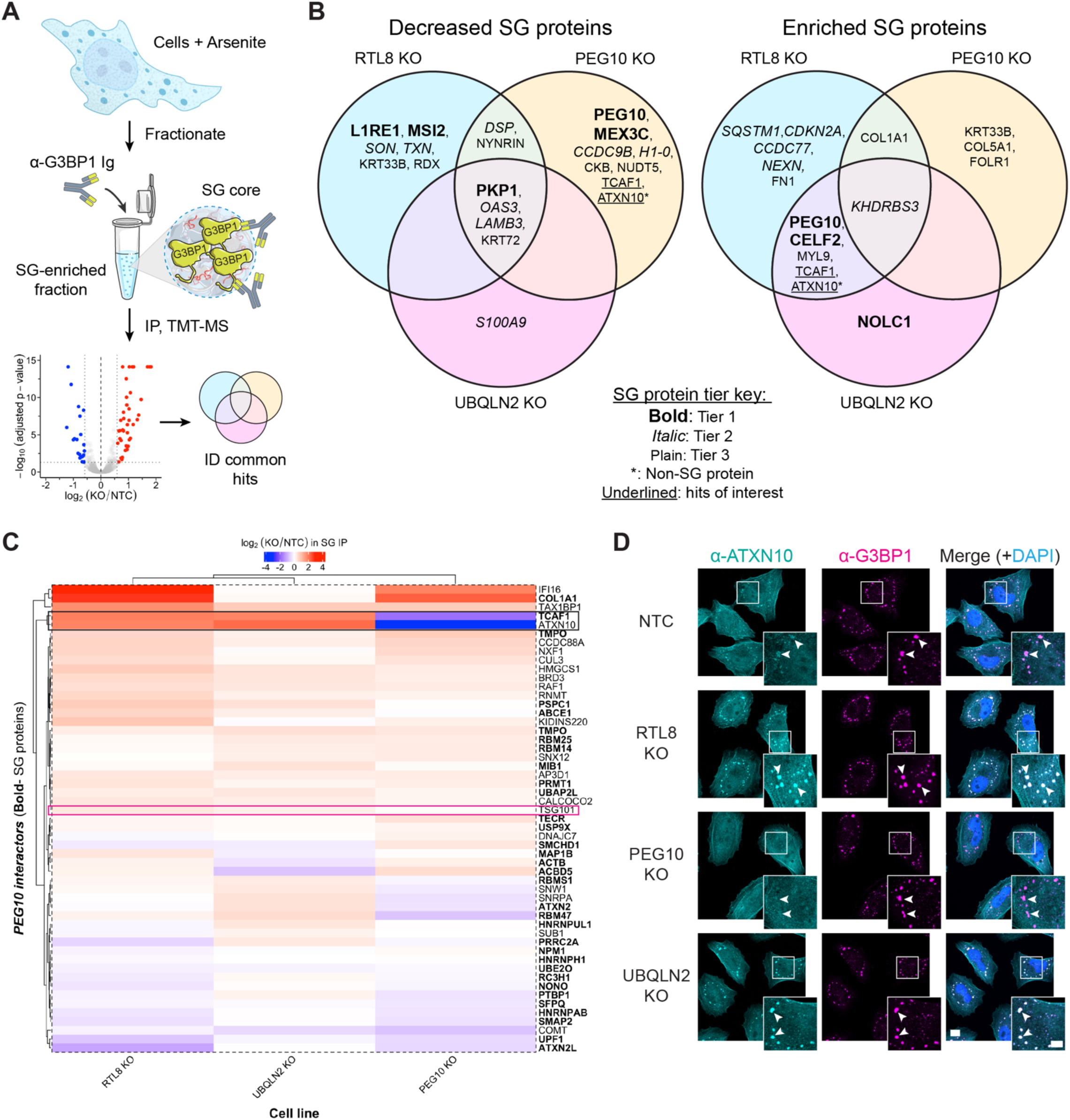
PEG10 shuttles select EV proteins to SGs. **A)** Schematic of SG pulldown experiments from HeLa cell lines. Also, see Figure S7A. **B)** Venn diagrams showing significantly decreased or enriched SG proteins in the indicated HeLa cell lines. SG protein tiers are annotated based on the RNA granule database. Also, see Figure S7B and Data S2. **C)** Heatmap of known PEG10 interactors detected in SG IP/MS experiments. Bolded proteins are also SG proteins. Colors correspond to fold enrichment in respective KO line versus NTC. Black box identifies TCAF1 and ATXN10, and pink box identifies TSG101, an EV marker. Also, see Figure S7C and S7D. **D)** After treatment with sodium arsenite, the indicated cell lines were stained for endogenous ATXN10 and G3BP1. Nuclei are labeled with DAPI and arrowheads in insets mark SGs. Scale bars: 10 µm and 5 µm for insets. Also, see Figure S7E.

Intriguingly, TCAF1, a tier 3 SG protein, showed a strong dependence on PEG10 levels. It was among the top enriched proteins in SGs from RTL8 and UBQLN2 KO cells, and, conversely, was among significantly depleted proteins in SGs from PEG10 KO cells. As TCAF1 interacts with PEG10,^21^ we assessed whether other known PEG10 interactors in our data set, which includes several SG proteins, showed a dependence on PEG10 for their SG incorporation. Only ATXN10, which is not known to be a SG protein, displayed the same trend as TCAF1 (Figure 6C). TCAF1 and ATXN10 are both known to be incorporated into EVs in a PEG10-dependent manner.^21^ To understand whether PEG10 enables the delivery of other EV proteins to SGs and to rule out potential EV contamination in our pulldown, we queried a panel of EV proteins, including the PEG10 interactor, TSG101.^82,99^ We observed no significant differences, implying that TCAF1 and ATXN10 belong to a select, small group of proteins that undergo PEG10-dependent localization to SGs (Figure S7C). While total levels of TCAF1 were unaltered, ATXN10 levels showed an increase in RTL8 KO cells compared to NTC cells (Figure S7D).

By IF, we validated our findings with ATXN10 which increasingly localized to SGs in RTL8 and UBQLN2 KO cells and no longer localized to SGs in PEG10 KO cells (Figure 6D). Like UBQLN2, ATXN10 did not localize to SGs in PEG10 gag-pol KO cells confirming that the gag-pol isoform mediates ATXN10 localization to SGs (Figure S7E). We could not carry out similar studies for TCAF1 due to a lack of suitable anti-TCAF1 antibodies for IF. Taken together, these data suggest that PEG10, RTL8, and UBQLN2 uniquely affect SG composition and that PEG10 shuttles select EV proteins to SGs.

### PEG10-derived VLPs are present within SGs

PEG10 is among a handful of capsid (CA)-like domain-containing mammalian genes that form VLPs, which can transport RNA and even proteins in EVs.^4,21,27–34^ The PEG10-dependent increase in EV proteins within SGs raises the intriguing possibility that PEG10 also exists as VLPs within SGs. To test this, we applied a custom cryo-correlative light and electron microscopy (cryo-CLEM) workflow^86^ to cells expressing PEG10 gag-pol-Dendra2 and harboring SGs labeled with G3BP1-mScarlet-I.

We first verified that PEG10 gag-pol-Dendra2 could still form VLPs by expressing it in PEG10 KO HeLa cells and concentrating VLPs from the media by ultracentrifugation (Figure 7A). Using a protease protection assay, we documented the presence of membrane-protected PEG10 gag-pol-Dendra2 in our VLP suspensions, which also contained known PEG10 interactors TCAF1, ATXN10, and RTL8^21^ (Figure S8A). We confirmed PEG10 oligomers by size exclusion chromatography and the presence of VLPs by negative stain transmission electron microscopy (TEM, Figure S8B and C). Subsequently, we applied the VLP suspension to Holey carbon-supported copper grids, plunge-froze them in liquid ethane, and used cryo-confocal microscopy to image regions positive for Dendra2 fluorescence on the grids. Enveloped VLPs, 50-100 nm in size, were detected, matching previous size estimates for PEG10 VLPs^21,29,31^ (Figure 7B and C). Thus, tagging PEG10 with Dendra2 still enables VLP formation.

**Figure 7:**
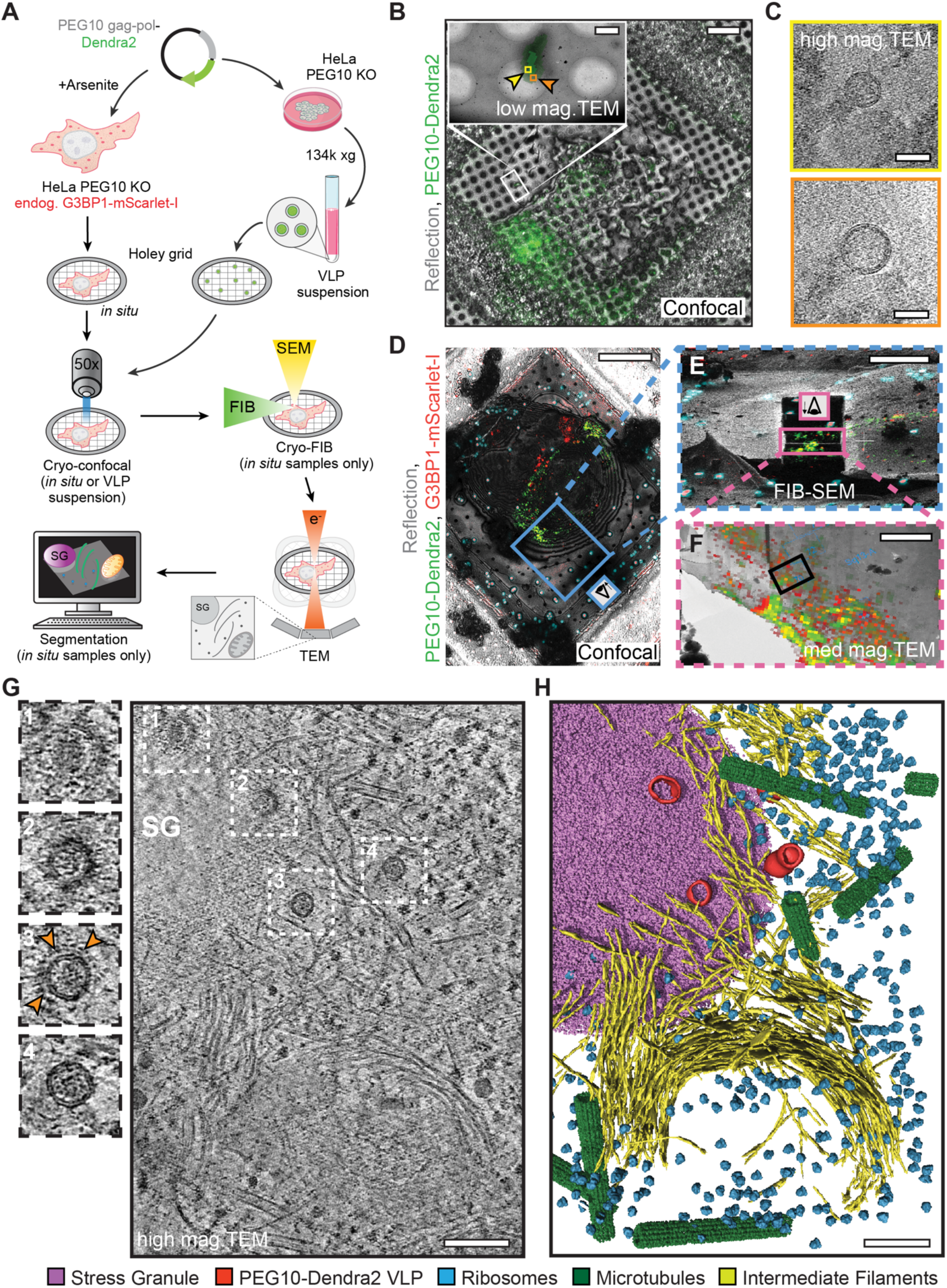
PEG10-derived VLPs are present within SGs. **A)** Schematic of the cryo-CLEM workflow for imaging PEG10 gag-pol-Dendra2-labeled VLPs in cells (i.e. *in* situ) or isolated from cell culture media (VLP suspension). **B)** Representative maximum intensity projection (MIP) of vitrified VLPs from PEG10 KO HeLa cells transfected with PEG10 gag-pol-Dendra2 overlaid on the reflection channel image. The inset depicts a low-magnification TEM image targeting the Dendra2 fluorescence signal. Also, see Figures S8A-C. Scale bars: 20 µm and 0.5 µm for overlay and inset images, respectively. **C)** Slice through a tomogram of the region highlighted in **B** showing two representative PEG10 gag-pol-Dendra2 VLPs. Scale bar: 50 nm. **D)** Representative overlay of MIP of cryo-confocal Z-stack with the image from the reflection channel (gray) of arsenite-treated vitrified PEG10 KO HeLa cells expressing endogenous G3BP1-mScarlet-I, transfected with PEG10 gag-pol-Dendra2. Fiducial fluorospheres (blue) were added for correlation. Scale bar: 20 µm. Also, see Figure S8D. **E)** Cryo-focused ion beam view post-milling with transformed cryo-confocal MIP matching the 15° milling angle. Scale bar: 10 µm. **F)** Overlay of medium magnification montage from cryo-TEM (6500x) of a lamella with cryo-confocal MIP. The solid black rectangle indicates where the tilt series was subsequently acquired. Scale bar: 2 µm. **G)** A slice through the tomogram of the region highlighted in F was collected at 42,000x, revealing well-preserved cellular interiors. Four (1-4) PEG10 gag-pol-Dendra2 VLPs in the field of view are highlighted in white dashed rectangles. Enlarged views of the four VLPs are also shown to highlight the variability in structure and morphology, such as additional protein coat (orange arrowheads, rectangle 3). Scale bar: 200 nm. Also, see Figure S8E. **H)** Segmentation of the tomogram volume shown in **G.** Stress granules (pink), PEG10 gag-pol-Dendra2 VLP (red), ribosomes (blue), intermediate filaments (yellow), and microtubules (green) are highlighted. Also, see Movie S3.

To investigate the architecture of PEG10 VLPs in the native cellular context (i.e. *in situ*), we performed cryo-CLEM on arsenite-stressed PEG10 KO HeLa cells expressing endogenous G3BP1-mScarlet-I and transfected with PEG10 gag-pol-Dendra2 at low expression levels mirroring that of endogenous gag-pol in RTL8 KO cells (Figure 7A and S8D). After identifying regions of mScarlet-I and Dendra2 colocalization by cryo-confocal microscopy (Figure 7D), we obtained thin lamellae of cells by cryo-focused ion beam (FIB) milling (Figure 7E and F). Subsequently, cryo-electron tomography of the correlated regions revealed well-preserved intact cellular structures, including SGs that are smooth in the center and grainy at the periphery, resembling other RNP condensates^100^ (Figure 7G). Remarkably, numerous VLPs localized within or adjacent to SGs, as highlighted in the segmentation of a reconstructed tomogram (Figure 7H and Movie S3). Observed VLPs were, on average, 67.3 nm in diameter (Figure S8E). In some instances, the VLPs were partially decorated, seen as projections on their surface (Figure 7G). In summary, our results show that PEG10-derived VLPs are present in SGs and highlight remarkable structural heterogeneity within SGs.

## DISCUSSION

The biological functions of endogenous retrovirus-like proteins are largely unknown. In this study we identified pivotal roles for a pair of such proteins in linking and modulating PQC and stress response pathways. The PQC factor UBQLN2 interacts with and regulates one retrovirus-like protein, PEG10, in a manner that requires a second, related retrovirus-like protein, RTL8. RTL8, in turn, enables PEG10 to shuttle UBQLN2 to SGs. The loss of any of these three proteins affects SG dynamics and composition. Our discovery of PEG10-derived VLPs in SGs also underscores the structural heterogeneity of SGs and raises the possibility of a virus-like mechanism for delivering molecules to or from SGs. Our results establish novel co-opted functions for retrovirus-like proteins in UBQLN2-mediated stress response and more broadly in SG biology.

UBQLN2 stabilizes RTL8 yet selectively promotes degradation of the gag-pol isoform of PEG10.^22,50,51^ A stable pool of RTL8 is required for PEG10 gag-pol turnover by UBQLN2, and other regulators of PEG10 (for example, the ubiquitin ligase UBE3A^21^) cannot compensate for the loss of RTL8. This dichotomous regulation of RTLs by UBQLN2, coupled with recent evidence that RTL8 inhibits PEG10 VLP production, suggest RTL8 has evolved to regulate PEG10 via UBQLN2. This notion is supported by phylogenetic data indicating that PEG10 and RTL8 arose from separate retrotransposon domestication events.^4^ Consistent with this view, the retrovirus-like protein most closely related to RTL8, LDOC1, also interacts with PEG10^16^ but cannot replace RTL8, implying a non-redundant role for RTL8 in PEG10 regulation. The unique N-terminus of RTL8, consisting of helical and disordered moieties, appears essential for PEG10 regulation. It mediates RTL8 binding to both UBQLN2 and PEG10, and so brings UBQLN2 and PEG10 together with two important consequences: it promotes PEG10 clearance while also permitting both UBQLN2 and RTL8 localization to SGs. The N-termini of RTL family members vary considerably and are hypothesized to confer novel functions to each protein.^4^ Our study presents evidence supporting this hypothesis. PEG10 may have also directed the evolution of RTL8’s N-terminus, suggested by the fact that rodent PEG10 and the N-termini of rodent RTL8 proteins are highly dissimilar compared to other mammals.^22,24,33,51^

Though our PLA results support continued interactions between UBQLN2 and PEG10 within SGs, super resolution microscopy indicates that they may localize to different subregions. PEG10 localizes to SG cores, whereas UBQLN2 appears to be excluded from the cores. These observations highlight heterogeneity in the distribution of PEG10 and UBQLN2 that may specify their functions within SGs and yields further support to the “core and shell” model of SG organization.^92,101^ Since UBQLN2 and RTL8 can phase-separate together^51^ and PEG10 is also predicted to phase-separate,^102^ further understanding how interactions between the three proteins shape their collective phase separation tendencies may shed light on the biophysical processes driving SG recruitment.

UBQLN2 is known to limit SG formation, and its depletion leads to increased cell death once stress resolves.^36,89^ The contribution of PEG10 to SG biology, however, remains an open question. In our study, PEG10-dependent delivery of UBQLN2 to SGs occurred with all tested stressors and in all tested cell lines, supporting a key role for PEG10 in SG biology. Though both PEG10 isoforms possess IDRs, only the gag-pol isoform selectively interacts with UBQLN2 and RTL8, undergoes B-isox-mediated precipitation, and localizes to SGs. Eliminating the disordered C-terminal PPR of gag-pol does not affect its localization to SGs, implying that other structural determinants within the pol region are responsible.^103^ Intriguingly, in the absence of PEG10 gag-pol alone, UBQLN2 is excluded from the nucleus (Figure S5B). Conceivably, PEG10 gag-pol or its cleavage products may traffic UBQLN2 to the nucleus, where UBQLN2’s presence is thought to be important during proteotoxic stress.^36,104^

We find that depletion of UBQLN2, RTL8, or PEG10 affects several aspects of SG dynamics and composition. By assessing endogenous G3BP1 labeled with a fluorescent tag, our analyses of SG dynamics avoid overexpression artifacts which can confound the study of biomolecular condensates.^56^ Depletion of UBQLN2 accelerates SG assembly, consistent with previous studies.^89^ Interestingly, SG formation is unchanged in RTL8 and PEG10 KO cells, both of which lose UBQLN2 localization to SGs. This result suggests that UBQLN2’s influence on SG formation occurs when it is present elsewhere in the cell rather than within the SGs themselves, indicating that UBQLN2 may act outside of SGs to limit their formation. Conversely, all three proteins appear to slow down SG dissolution. Furthermore, PEG10 KO cells as well as RTL8 and UBQLN2 KO cells, which express more PEG10, disassemble SGs faster. Therefore, it is reasonable to conclude that an optimal amount of PEG10 is needed for proper SG dissolution. In addition to changes in dynamics, we find significant changes in SG size in our KO lines. However, the exact role of UBQLN2, RTL8, and PEG10 in regulating SG size remains to be studied.

Our proteomic analyses of SG composition reveal compelling evidence for PEG10-mediated delivery of two EV proteins, ATXN10 and TCAF1,^21^ to SGs. Our studies establish ATXN10 as a second novel SG protein (the first being RTL8) whose incorporation into SGs is dependent on PEG10 levels. Since the gag-pol isoform is necessary for ATXN10 delivery to SGs, we speculate that it also delivers TCAF1 to SGs. ATXN10 activates MAPK signaling, whereas TCAF1 participates in calcium signaling during stress.^60,105^ SG sequestration of MAPK activators, for example PKC, has been shown to inhibit MAPK activation and reduce cell death.^106,107^ Therefore, by sequestering ATXN10 and TCAF1 in SGs, PEG10 may regulate downstream signaling pathways, such as MAPK and calcium signaling, and ultimately regulate processes like apoptosis. PEG10 may serve as a selective shuttle for proteins to SGs - thus far, we have identified four of its targets (UBQLN2, RTL8, ATXN10, and TCAF1), but there may be additional targets expressed early in development or disease. Disease-induced upregulation of PEG10 could also trigger the localization of these target proteins to pathological RNP condensates. To better understand these effects, it would be helpful to express PEG10 in naturally PEG10-deficient cell models such as U-2 OS cells and certain neuronal subtypes. Furthermore, generating conditional knockout animal models of PEG10 would allow for an in-depth study of its functions under disease conditions.

While most of the changes we observe in SG composition are unique to each KO line, changes to proteins associated with viral stress response (MEX3C and OAS3) or originating from retroelements (L1RE1 and NYNRIN) are shared between RTL8 and PEG10 KO lines. SG formation is intimately connected to retroelement activation and is a means to defend against viral infections.^108–112^ Therefore, understanding how RTLs affect these pathways could uncover functionality beyond the scope of this study.

PEG10 belongs to a rarified group of retrovirus-like proteins that can form VLPs.^4,21,22,27–33^ Our cryo-CLEM studies reveal the presence of intact PEG10-derived VLPs in SGs, which underscores the structural heterogeneity of SGs. Far from being homogenous condensates, SGs are likely complex and possess local regions of demixing giving rise to oligomers and aggregates.^113^ Our results identify yet another level of complexity: VLPs embedded within SGs. Further work will be needed to clarify whether VLPs in SGs originate outside of SGs and are transported to them or arise within SGs due to spontaneous PEG10 oligomerization in demixed regions. VLPs are typically secreted in EVs^29^ and transport RNA between cells but have also been engineered to deliver artificial RNA and polypeptides.^30,31,34^ Because three of the four proteins we discovered to undergo PEG10-mediated delivery to SGs are either associated with EVs (ATXN10 and TCAF1) or can be incorporated into PEG10 VLPs (RTL8),^21,33^ it is conceivable that PEG10 VLPs serve to shuttle such proteins to SGs. These proteins may be present within VLPs or bound to their surface. Indeed, a significant fraction of VLPs we observe *in situ* have decorations that could be proteinaceous. Given that both PEG10 isoforms can generate VLPs^22^ yet only gag-pol localizes to SGs, VLP-forming ability alone is not sufficient to ensure SG delivery; gag-pol may possess as yet unknown determinants that “license” its VLPs to SGs. PEG10 VLPs could also deliver specific RNA cargo to SGs or extract RNA and/or proteins from SGs to aid in SG dissolution. Consistent with this latter possibility, increased PEG10 levels results in faster SG clearance.

Collectively, our study identifies critical functions for retrovirus-like proteins in stress response, namely the stress-induced actions of UBQLN2 and, more broadly, SG dynamics, composition and heterogeneity. Well-established examples of retrovirus-like proteins participating in mammalian physiology and disease are Arc and PNMA2.^6–8,27,28,32^ Our work expands the co-opted functions of retrovirus-like proteins to cellular stress pathways. While important for placental development,^11^ PEG10 is aberrantly increased in malignancies such as pancreatic cancer and hepatocellular carcinoma, and upregulation of gag-pol in particular is implicated in ALS and AS.^15,21,22,25,26^ Understanding the effects of PEG10 dysregulation on SG biology should provide mechanistic insight into these diseases, as activation of stress pathways is implicated in their pathology.^53,54,57–59,114–117^ Our findings also elevate RTLs from mere interactors with UBQLN2 into key contributors to UBQLN2 biology. UBQLN2 missense mutations that cause hereditary ALS are known to alter interactions with SG components.^98^ Determining whether retrovirus-like proteins, whose levels may change as ALS progresses, influence the pathological effects of UBQLN2 mutations will be an important next step.

### Limitations of the study

The data generated in this study were obtained using endogenous levels of proteins in immortalized cells. To understand implications for disease, disease-relevant systems such as iPSC-derived neurons and rodent models will be important. SGs are an acute response to stress and may not faithfully recapitulate the chronic low-level stress that occurs during disease.^118^ Therefore, future studies should seek to understand how retrovirus-like proteins and UBQLN2 affect chronic stress responses. We find strong PEG10-dependent localization of ATXN10 and TCAF1 to SGs, but it is possible that subsequent studies employing higher sensitivity proteomics may uncover more such proteins. Lastly, our *in situ* cryo-CLEM experiments were achieved by overexpressing PEG10 in PEG10-deficient cells, which raises the possibility that the occurrence of VLPs in SGs is not a widespread phenomenon. VLPs may also disassemble when they are in SGs, further complicating their detection. Though technically challenging, additional experiments

would be needed to determine whether PEG10 VLPs are present in SGs at endogenous levels of expression and whether perturbations to overall PEG10 levels alter the occurrence of VLPs in SGs.

## Author contributions

Conceptualization, H.M.M., A.M.W., S.J.B., S.M., H.L.P., and L.M.S.; Methodology, H.M.M., M.G.F., C.H., V.B., S.M., H.L.P., and L.M.S.; Software, H.M.M., C.H., and D.S.; Investigation, H.M.M., M.G.F., C.H., R.L., J.H.R., and N.G.; Writing – Original Draft, H.M.M., H.L.P., and L.M.S.;

Writing – Review and Editing, H.M.M., A.M.W., S.J.B., S.M., H.L.P., and L.M.S.; Funding Acquisition, S.J.B., S.M., H.L.P., and L.M.S.; Resources, A.M.W. and S.J.B.; Supervision, V.B., S.M., H.L.P., and L.M.S.

## Acknowledgments

We thank Zhongsheng You for the TCAF1 antibody, Peter Howley for the FLAG-UBQLN2 plasmid, Feng Zhang for the PX459 plasmid, and Jürgen Knoblich for the G3BP1-mScarlet-I HDR plasmid. We are grateful to Carlos Castañeda and Thuy Dao for their valuable feedback and guidance, and to Stephanie Moon and Benjamin Dodd for their advice on CRISPR-tagging G3BP1 and live-cell imaging. We thank the University of Michigan Microscopy Core and Proteomics Resource Facility for their assistance with imaging and proteomics, respectively. We also acknowledge all members of the Paulson, Sharkey, Barmada, Todd and Srinivasan labs for their suggestions, camaraderie, and sharing of various reagents. Third-party graphical elements were incorporated from Bioicons under a CC-0 license, Servier under CC-BY 3.0 Unported license, and DBCLS under CC-BY 4.0 Unported license. Much of the research described in this study was conducted in the HeLa cell line, which was established from Henrietta Lacks’ tumor cells without her knowledge or consent in 1951. We are grateful to Henrietta Lacks, now deceased, and to her surviving family members for their contributions to biomedical research. This work was supported by National Institutes of Health (T32GM141840 to M.G.F.; T32GM007863 to N.G.; R01NS097542, R01NS113943, and 1R56NS128110-01 to S.J.B; DP2GM150019-01 to S.M., and R35NS122302 and P30AG072931 to H.L.P.), Rackham Graduate School (Predoctoral Fellowship to H.M.M.), Lefkofsky Family Foundation (early-career award to L.M.S.), and Klatskin Sutker Discovery Fund (to S.M.). The University of Michigan cryo-EM facility is supported by NIH S10OD030275 and the Arnold and Mabel Beckmann Foundation award.

---

**Supplementary Figure 1:**
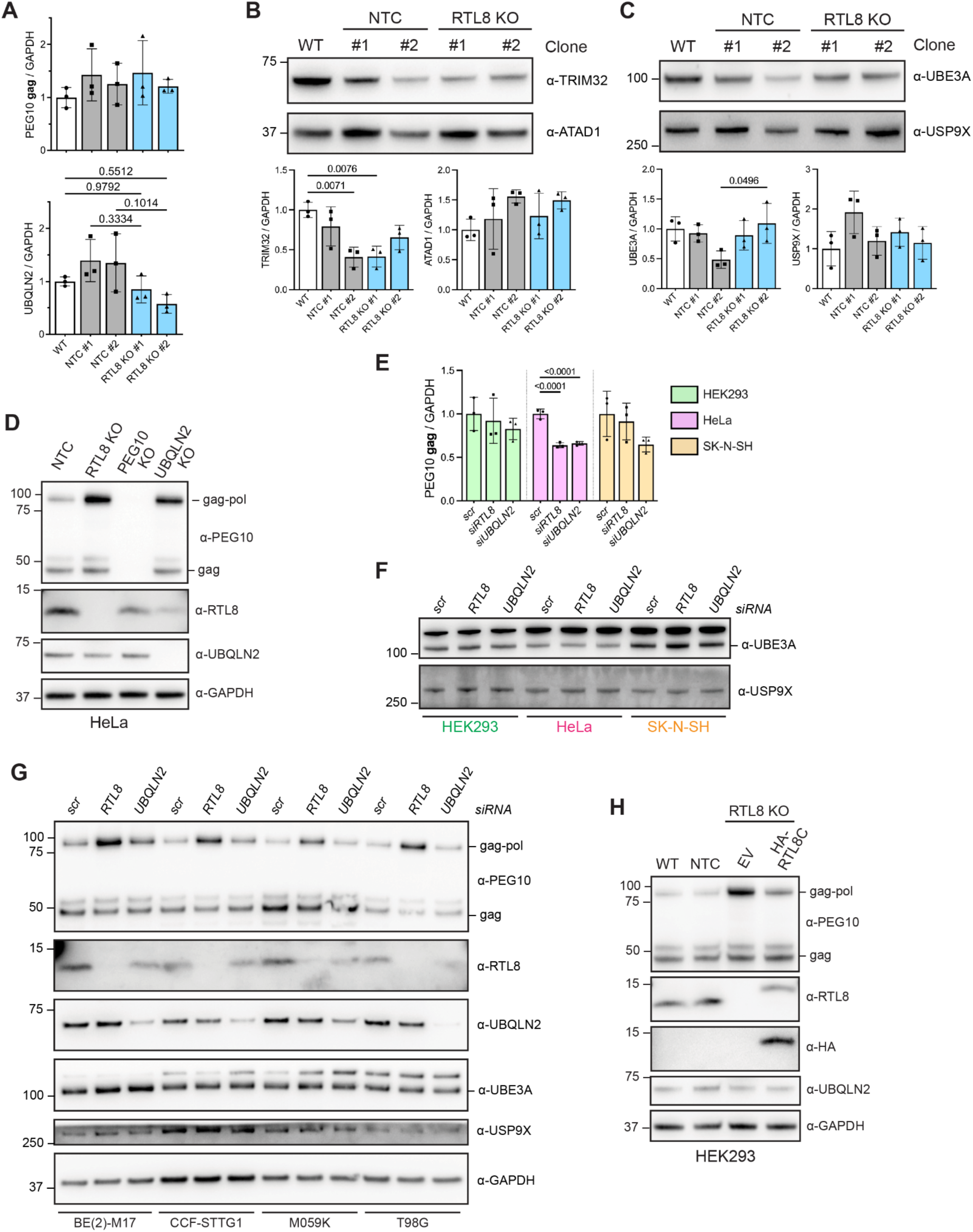
RTL8 depletion selectively increases PEG10 gag-pol levels and rescuing RTL8 is sufficient to restore PEG10 levels, related to Figure 1. **A)** Relative UBQLN2 and PEG10 gag levels in lysates from WT, NTC and RTL8 KO HEK293 cells, shown in Figure 1C. **B)** Immunoblot of lysates from WT, NTC and RTL8 KO HeLa cells probed for UBQLN2 client proteins, TRIM32 and ATAD1. Relative levels of TRIM32 and ATAD1, normalized to levels in WT cells, are shown below. **C)** Immunoblot of lysates from WT, NTC and RTL8 KO HeLa cells probed for known PEG10 regulators, UBE3A and USP9X. Relative levels of UBE3A and USP9X, normalized to levels in WT cells, are shown below. **D)** Immunoblot of lysates from NTC, RTL8 KO, PEG10 KO and UBQLN2 KO HeLa cell lines, probed for PEG10, RTL8 and UBQLN2. **E)** Relative PEG10 gag levels in different human cell lines following knockdown of RTL8 or UBQLN2, normalized to levels in control siRNA-treated cells, shown in Figure 1F. **F)** Immunoblot of lysates from different human cell lines probed for UBE3A and USP9X following knockdown of RTL8 or UBQLN2. **G)** Immunoblot of lysates from multiple human CNS-derived cell lines treated with siRNA against RTL8 or UBQLN2. **H)** Immunoblot of lysates from WT, NTC, and RTL8 KO HeLa cells stably expressing HA-RTL8C or empty vector. In (A-C & E), data represent means ± SD (N=3), analyzed with one-way ANOVA with Tukey’s multiple comparison test. GAPDH = loading control.

**Supplementary Figure 2:**
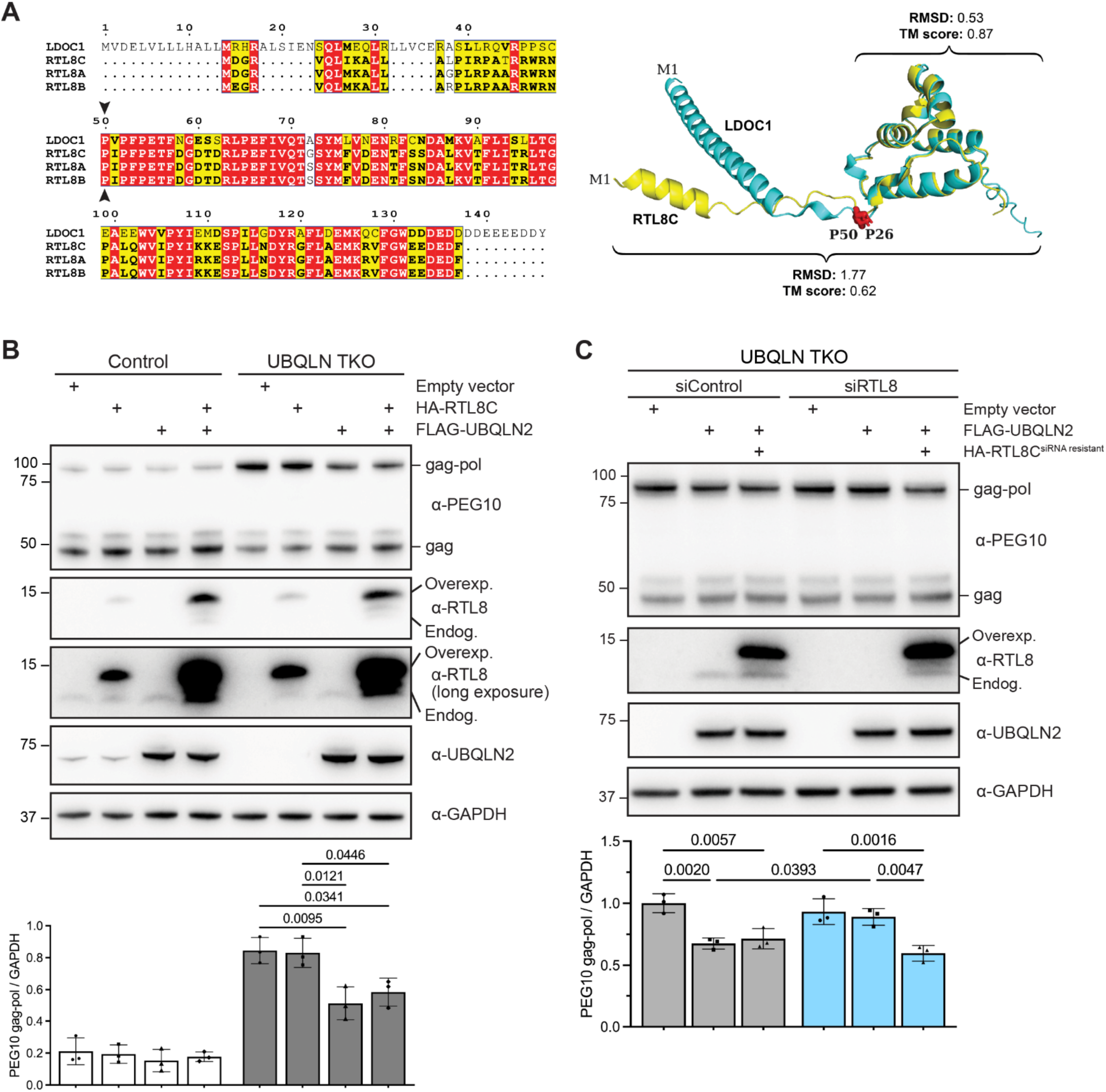
RTL8 acts on PEG10 gag-pol in a UBQLN2-dependent manner, related to Figure 1. **A)** At left, multiple sequence alignment between LDOC1 and RTL8 paralogs. Identical residues are boxed in red, similar residues are boxed in yellow and bolded residues are found in the majority of proteins. Arrowheads indicate Pro50 in LDOC1 (Pro26 in RTL8). At right, structure alignment between AlphaFold-predicted models of LDOC1 (cyan) and RTL8C (yellow). M1 denotes N-terminal methionine. Pro50 in LDOC1 and Pro26 in RTL8C are shown as red stick residues. Root mean square deviation (RMSD) values and template modeling (TM) scores for the full-length proteins and residues after Pro50 in LDOC1 (Pro26 in RTL8C) are shown above and below the alignment, respectively. **B)** Immunoblot of lysates from T-rex control or UBQLN TKO HEK293 cells expressing HA-RTL8C, FLAG-UBQLN2, both, or empty vector. Relative PEG10 gag-pol levels are shown below the blots. **C)** Immunoblot analysis of lysates from UBQLN TKO HEK293 cells treated with either control or RTL8 siRNA. Cells were transfected with either an empty vector or FLAG-UBQLN2 with or without co-expression of siRNA-resistant HA-RTL8C. Relative PEG10 gag-pol levels, normalized to cells transfected with control siRNA and empty vector, are quantified and shown below the blots. In (B & C), data represent means ± SD (N=3), analyzed with one-way ANOVA with Tukey’s multiple comparison test. GAPDH = loading control.

**Supplementary Figure 3:**
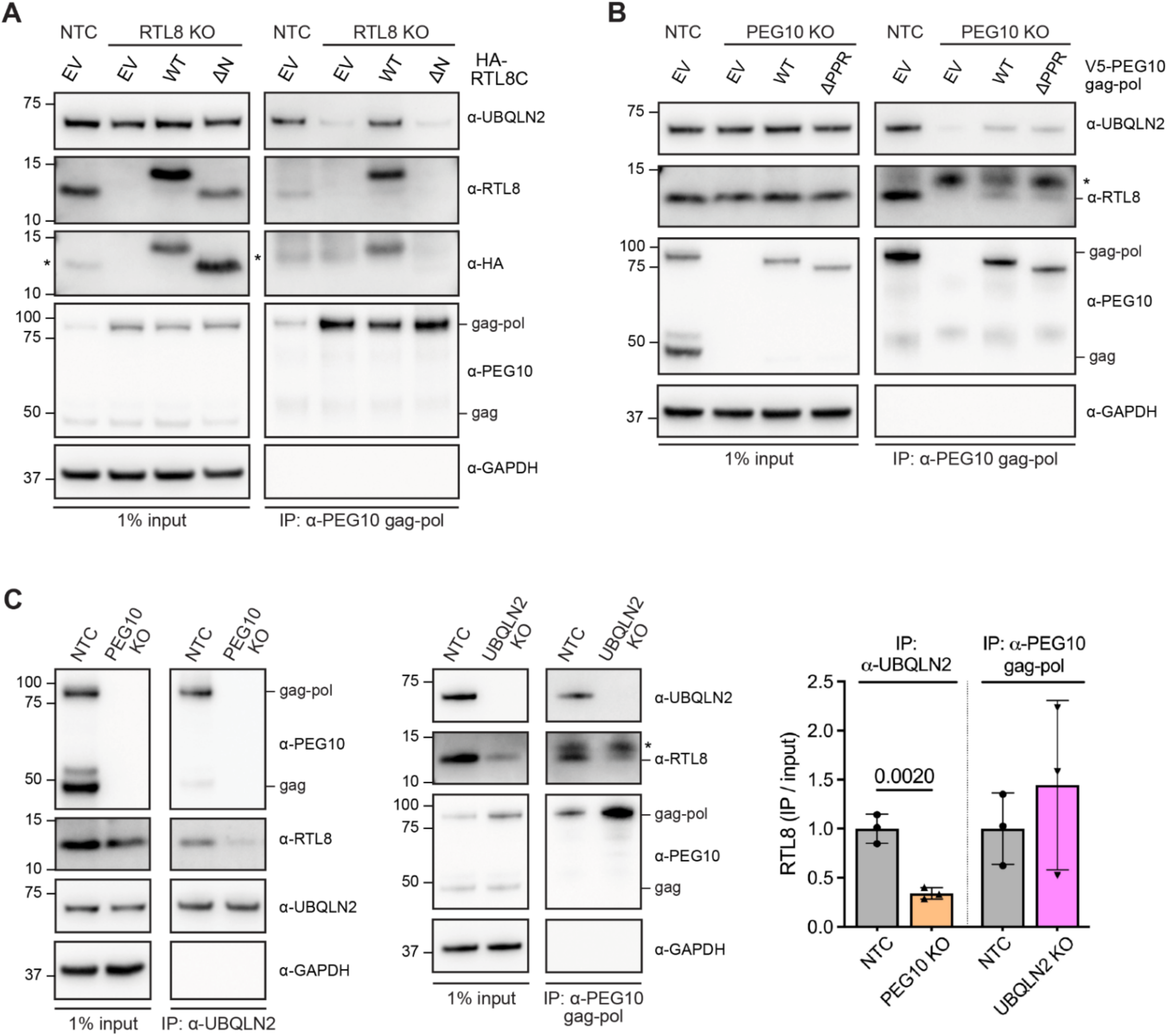
PEG10-UBQLN2 interactions depend on the N-terminus of RTL8 but not the PPR of PEG10, with evidence suggesting that RTL8 and PEG10 interact first. Related to Figure 2. Lysates from **A)** NTC or RTL8 KO HeLa cells transfected with either an empty vector or a vector expressing HA-RTL8C or HA-RTL8C ΔN, and **B)** NTC or PEG10 KO HeLa cells transfected with an empty vector or vector expressing V5-PEG10 gag-pol or V5-PEG10 gag-pol ΔPPR were subjected to IP using an α-PEG10 gag-pol antibody. **C)** Lysates from NTC and PEG10 KO HeLa cells IPed with an α-UBQLN2 antibody are shown on the right. Lysates from NTC and UBQLN2 KO HeLa cells IPed with α-PEG10 gag-pol antibody are shown in the middle. The amount of RTL8 co-IPed, relative to input and normalized to NTC HeLa within each co-IP, is shown on the left. * denotes non-specific band.

**Supplementary Figure 4:**
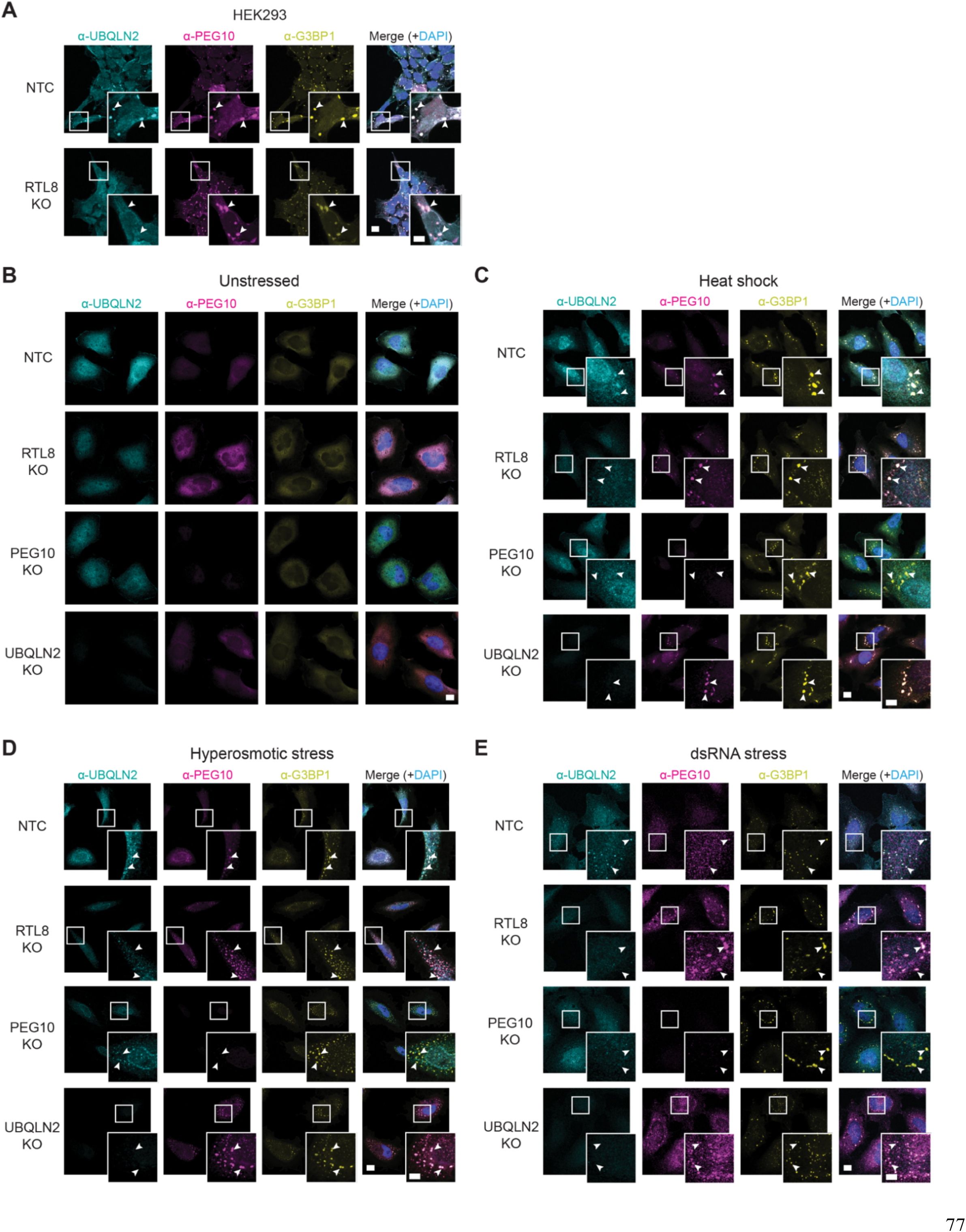
PEG10-mediated recruitment of UBQLN2 to SGs occurs in all tested stress conditions and cell lines, related to Figure 4. **A)** NTC and RTL8 KO HEK293 cells subjected to arsenite stress. NTC, RTL8 KO, PEG10 KO and UBQLN2 KO HeLa cells subjected to **B)** no stress, **C)** heat shock, **D)** hyperosmotic stress or **E)** dsRNA stress. In (A-E), cells are stained for endogenous UBQLN2, PEG10, and G3BP1. Nuclei are labeled with DAPI and arrowheads in insets mark SGs. Scale bars: 10 µm and 5 µm for insets.

**Supplementary Figure 5:**
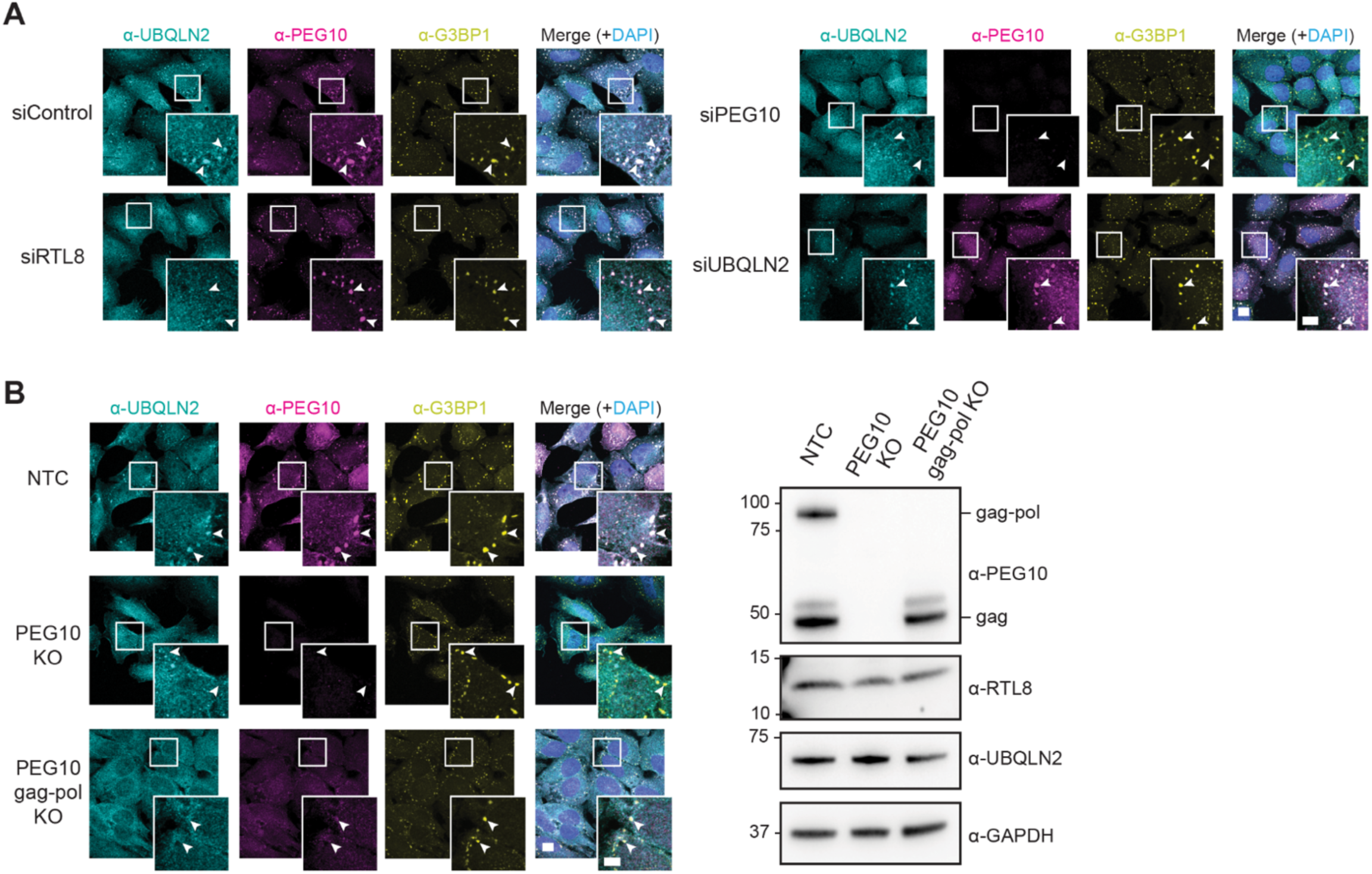
Depletion of PEG10 gag-pol prevents UBQLN2 localization to SGs, related to Figure 4. **A)** WT HeLa cells treated with non-targeting control siRNA or siRNA against RTL8, PEG10 or UBQLN2, then treated with sodium arsenite. **B)** NTC, PEG10 KO and PEG10 gag-pol KO HeLa cells treated with sodium arsenite are shown on the right. Immunoblot of lysates from respective cell lines probed for the indicated proteins are shown on the left. GAPDH = loading control. In (A & B), cells are stained for endogenous UBQLN2, PEG10, and G3BP1. Nuclei are labeled with DAPI and arrowheads in insets mark SGs. Scale bars: 10 µm and 5 µm for insets.

**Supplementary Figure 6:**
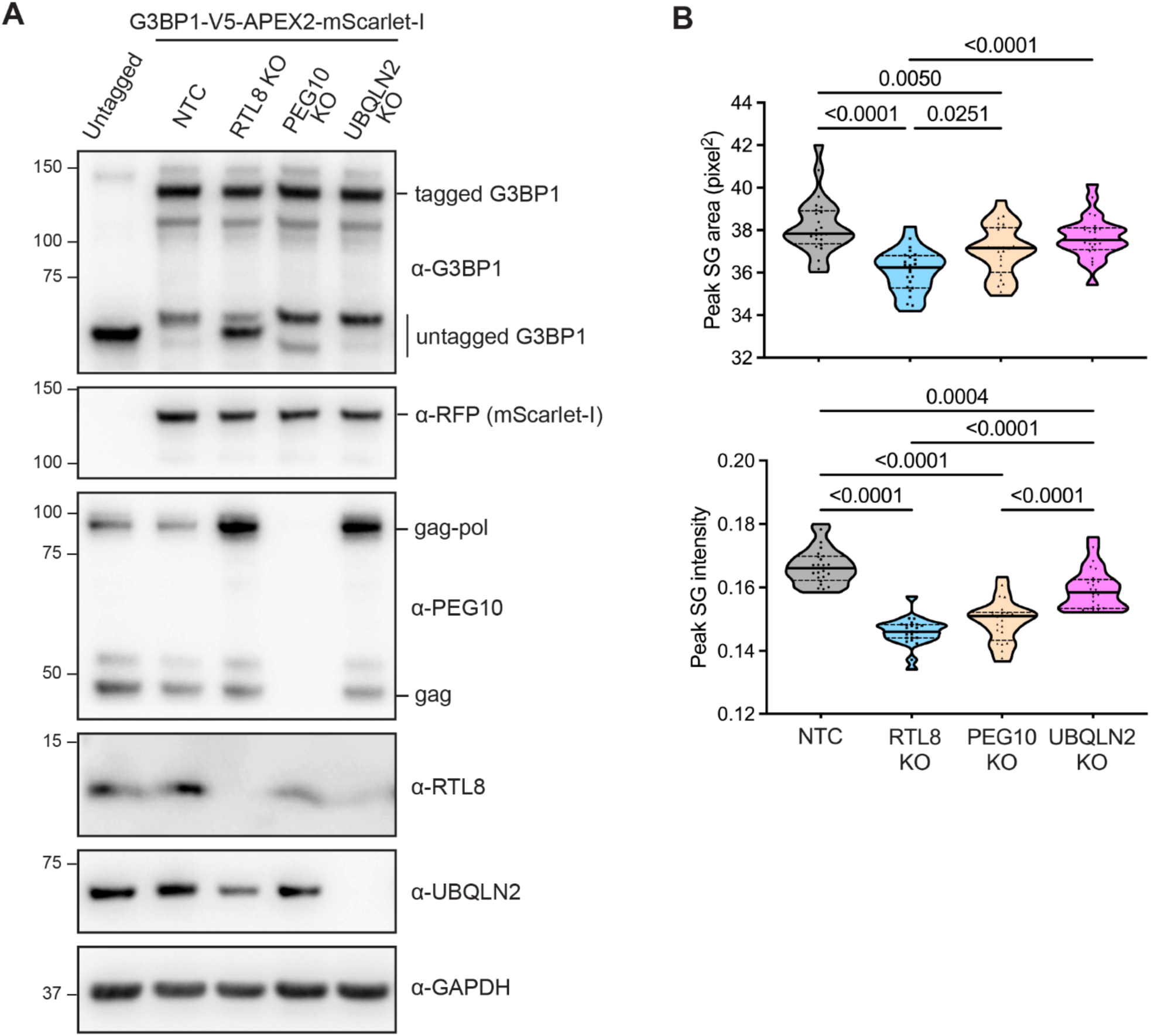
Depletion of UBQLN2, RTL8, or PEG10 results in changes in SG size, related to Figure 5. **A)** Immunoblot of lysates from the indicated cell lines, where endogenous G3BP1 has been CRISPR-tagged with V5-APEX2-mScarlet-I. Lysate from untagged NTC HeLa cells is also included for comparison. GAPDH = loading control. **B)** Area and intensity of SGs from the indicated cell lines in Figure 5A after 20 minutes of arsenite treatment as shown in Figure 7B. A one-way ANOVA with Tukey’s multiple comparison test was performed. N= 24 fields across 3 independent replicates.

**Supplementary Figure 7:**
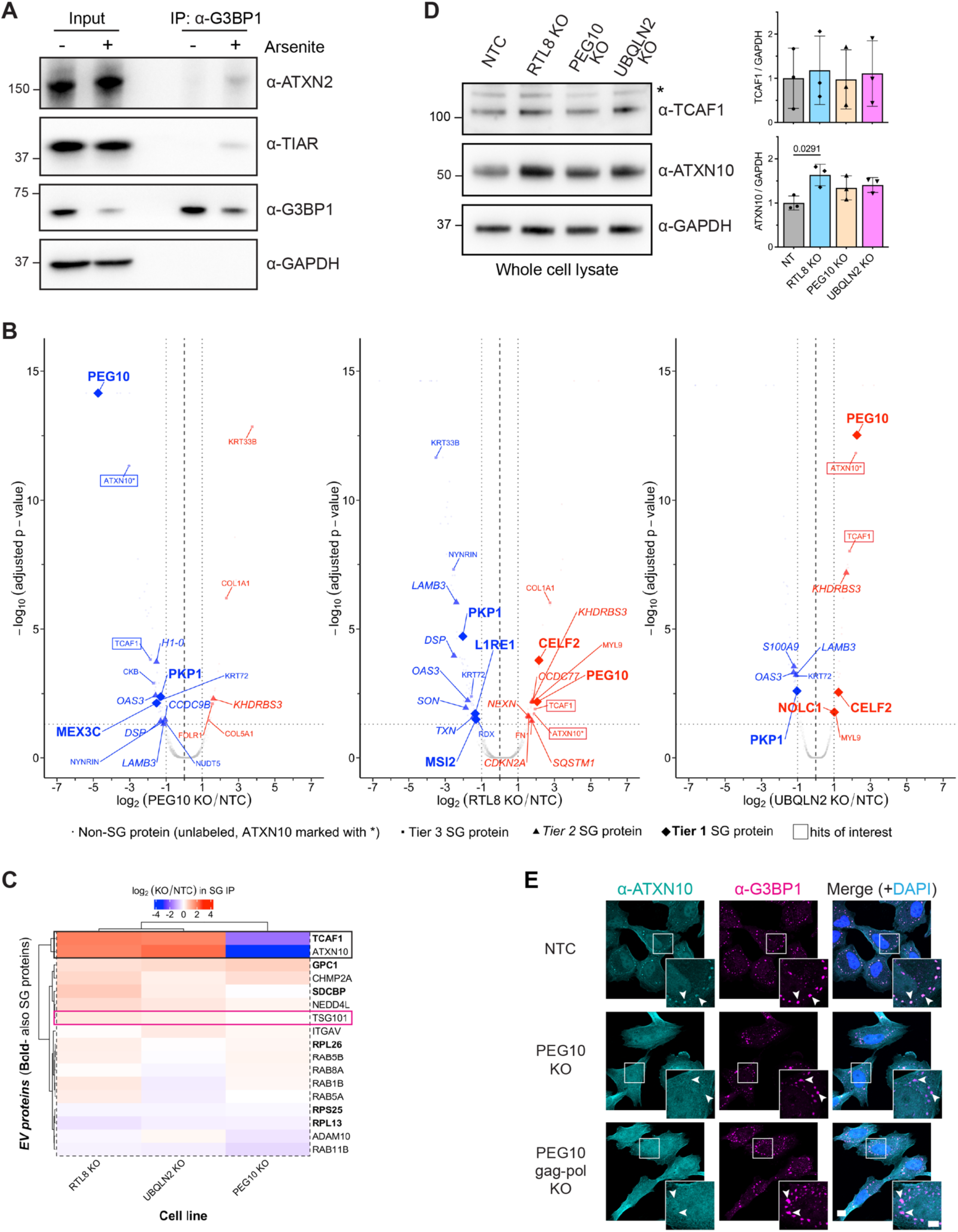
Analysis of SG IP results and evidence that the gag-pol isoform of PEG10 selectively shuttles TCAF1 and ATXN10 to SGs, related to Figure 6. **A)** Immunoblot of lysates from untreated and arsenite-treated HeLa cells subjected to SG pulldown with ⍺-G3BP1 Ig. ATXN2 and TIAR are positive controls, GAPDH is a negative control. **B)** Volcano plot of proteins significantly increased (red) or decreased (blue) in the indicated KO HeLa line compared to NTC cells. N = 4 biological replicates. The cutoff was set at p <0.05 and absolute fold change of 2. Labels and symbols are based on annotations of SG protein tiers on the RNA granule database. Boxes identify TCAF1 and ATXN10. **C)** Heat map of commonly used EV markers detected in our SG IP/MS experiments. Bolded hits are also SG proteins. Colors correspond to fold enrichment in respective KO line versus NTC. Black box identifies TCAF1 and ATXN10 and pink box identifies the PEG10 interactor, TSG101. **D)** At left, immunoblots of lysates from the indicated cell lines probed for TCAF1 and ATXN10. At right, relative TCAF1 and ATXN10 levels in the indicated lines, normalized to levels in NTC cells. Data represent means ± SD (N=3), analyzed with one-way ANOVA with Tukey’s multiple comparison test. GAPDH = loading control, * denotes non-specific band. **E)** Sodium arsenite-treated NTC, PEG10 KO and PEG10 gag-pol KO HeLa cells were stained for endogenous ATXN10 and G3BP1. Nuclei are labeled with DAPI and arrowheads in insets mark SGs. Scale bars: 10 µm and 5 µm for insets.

**Supplementary Figure 8:**
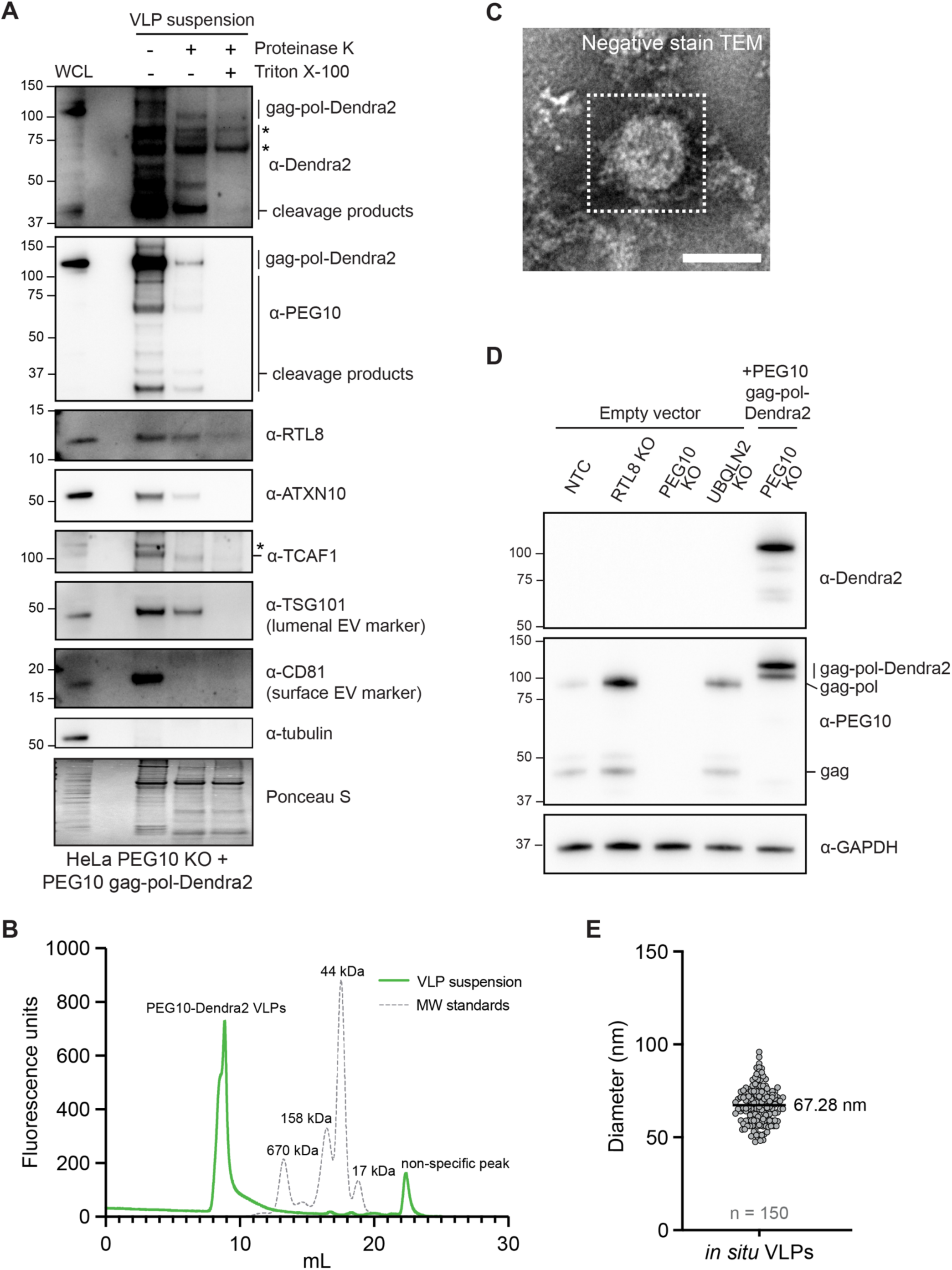
PEG10 gag-pol-Dendra2 forms membrane-enclosed VLPs, related to Figure 7. **A)** Immunoblot depicting a protease protection assay on PEG10 gag-pol-Dendra2 VLP suspensions. TSG101 and CD81 are EV markers, tubulin is used to show lack of cytoplasmic contamination, and Ponceau S is used to normalize loading. * denotes non-specific band. **B)** Fluorimeter trace of PEG10 gag-pol-Dendra2 VLP suspensions monitored via size-exclusion chromatography. **C)** Negative-stain TEM image of crude PEG10 gag-pol-Dendra2 VLP suspensions. White dashed box highlights a VLP. Scale bar: 50 nm. **D)** Immunoblot of lysates from indicated cell lines transfected with an empty vector or PEG10 KO cells transfected with PEG10 gag-pol-Dendra2. GAPDH = loading control. **E)** Size distribution of *in situ* PEG10 gag-pol Dendra2-VLPs. Mean diameter is indicated on the graph. n=150 VLPs.

**Data S1**: Proteomic analysis of RTL8 KO versus WT and NTC HEK293 cells, related to Figure 1.

**Data S2**: Proteomic analysis of SG cores pulled down from RTL8, PEG10 and UBQLN2 KO versus NTC HeLa cells, related to Figure 6.

**Movie S1**: Assembly of SGs in NTC, RTL8, PEG10 and UBQLN2 KO HeLa cells after arsenite addition, related to Figure 5.

**Movie S2**: Disassembly of SGs in NTC, RTL8, PEG10 and UBQLN2 KO HeLa cells during recovery from arsenite stress, related to Figure 5.

**Movie S3**: Segmentation of the *in situ* cryo-CLEM tomogram volume, related to Figure 7.

